# Improved spike-in normalization clarifies the relationship between active histone modifications and transcription

**DOI:** 10.1101/2025.11.25.690627

**Authors:** Lauren Patel, Yuwei Cao, Tianyao Xu, Eduardo Modolo, Tamar Dishon, Lingzhi Zhang, Eric Mendenhall, Sven Heinz, Itamar Simon, Christopher Benner, Alon Goren

## Abstract

Spike-in normalization enables quantitative analysis of ChIP-sequencing (ChIP-seq) signal. Here we introduce a novel robust dual-spike-in normalization approach for ChIP-seq (ChIP-wrangler). We identify optimal conditions, such as the ratio between the spike-in species and the target, demonstrate the ability of this approach to detect technical artefacts, and use ChIP-wrangler to revisit recent claims that active histone marks are dependent on transcription. Concerned that previous studies improperly used spike-in normalization to arrive at their conclusions, we used ChIP-wrangler to show that acute depletion of RNA polymerase II (RNAPII) has only a modest impact on the levels of H3K4me3 and H3K27ac. In line with other studies, our results provide proof that the maintenance of histone acetylation is not merely a consequence of ongoing transcription. Further, we show that promoters and enhancers are differentially impacted by inhibiting transcription. Specifically, of the 5.9% peaks that showed a decrease in H3K27ac following depletion of RNAPII, 82% are promoter-distal and contain enhancer-related DNA binding motifs. Further, the small subset of regions that gain acetylation (0.35%) were enriched for stress response motifs. Our innovative ChIP-seq normalization approach provides increased rigor and “guardrails” for successful spike-in normalization, and as applied here refines the understanding of the intricate crosstalk between RNAPII activity and histone marks associated with transcription.

## Main

A key step in quantitative analysis of ChIP-sequencing (ChIP-seq) data is normalization across samples or conditions. This is widely done using read depth normalization, but can be improved by adding exogenous spike-in chromatin to each sample as a reference. Spike-in normalization is a powerful approach to detect global changes in enrichment that read depth normalization may fail to quantify^1–3^. Yet, improper use of spike-in normalization can lead to artefacts and erroneous biological interpretations^4^. One key reason is that spike-in normalization relies on the ratio of spike-in to target chromatin reads, and mis-quantification of this single scaling ratio will skew the processed data which can have a profound impact on the interpretation of the results.

Here we present ChIP-wrangler, a robust normalization approach for ChIP-seq that employs spike-in chromatin from two species at a known ratio, to serve as a failsafe against possible technical errors. The dual spike-ins help track technical variability between samples, increase the rigor of quality control (QC) and thus minimize the risk of arriving at incorrect biological conclusions (**Figs. 1-2**). In the current implementation, we use *S. cerevisiae* and *D. melanogaster* chromatin (**Methods**), taking advantage of their distinct properties, such as genome size and average GC content of their genome, to allow parallel normalization and calculate confidence intervals for quantifying the target human ChIP-seq signal (**Figs. 1b-d, Supp. Figs. 1-2**; detailed experimental protocols and analysis framework are available in **Methods** and in a **GitHub repository**).

**Fig. 1.**
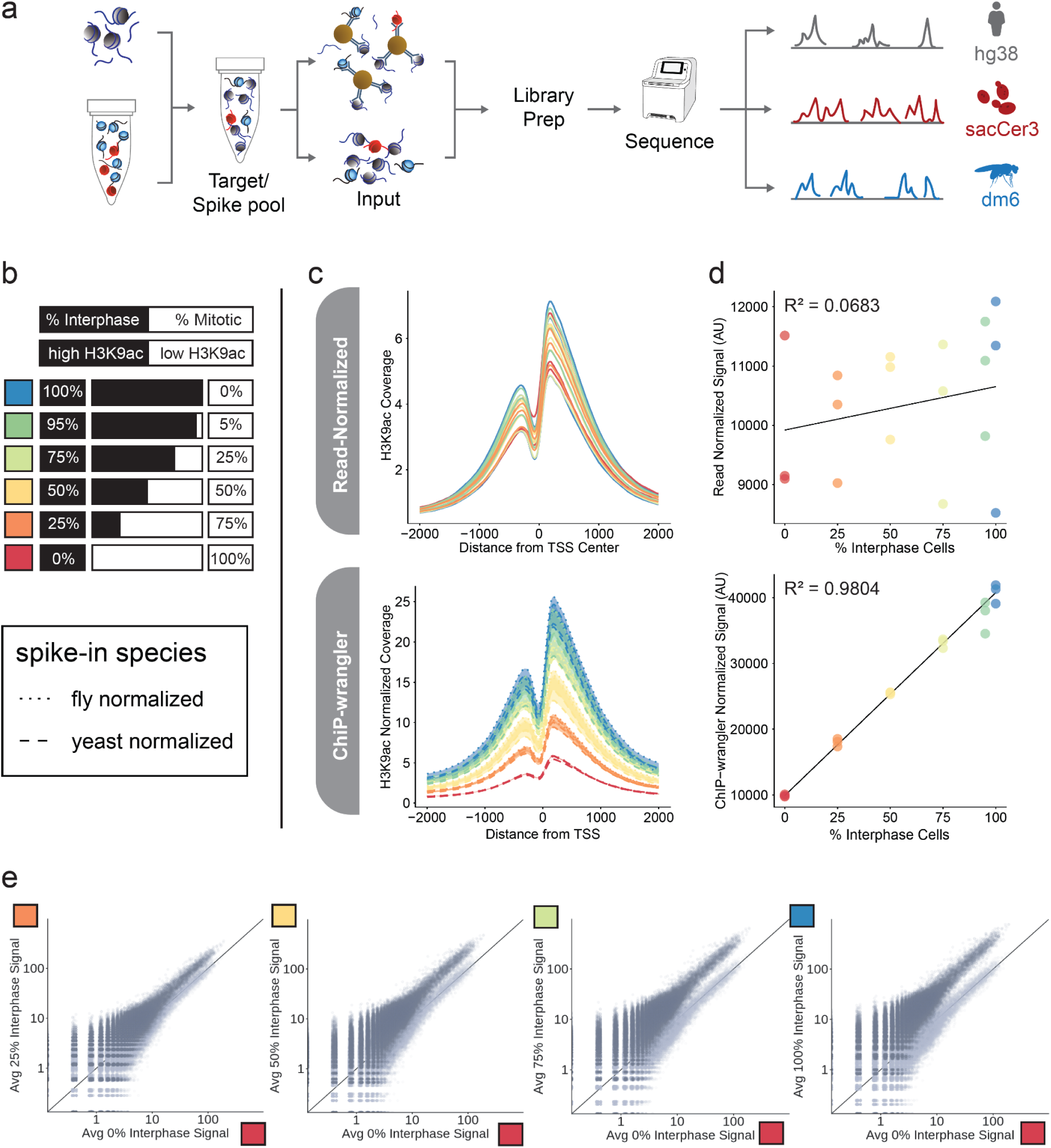
ChIP-wrangler enables rigorous quantification of ChIP-seq signal. **a.** Overview of the ChIP-wrangler experimental pipeline. Chromatin from two exogenous species (red and blue nucleosomes) is pre-mixed to minimize variability and then added to target chromatin (grey nucleosomes). Before immunoprecipitation, a subset of mixed chromatin is set aside to be used as an input control, from which the relative ratios of the exogenous species can be determined for spike-in normalization. The data is evaluated and analyzed as described in **Methods** to accurately quantify the signal in each species. **b.** A titration of H3K9ac signal was generated by mixing cells with low H3K9ac signal (mitotic-arrested HeLa-S3; white) with cells with high H3K9ac signal (unsynchronized HeLa-S3; black). The ratios are depicted as percentages in the bar graph. **c.** Metagene plot of H3K9ac at *H. sapiens* RefSeq promoters (TSS +/- 2kb) of read-depth normalized data (top) versus ChIP-wrangler normalized data (bottom). ChIP-wrangler’s confidence intervals (shaded bands) demonstrate variability between results of the separate normalization to each spike-in species. **d.** The area under the curve in (c) is plotted relative to the percent of interphase cells in each condition. The line connects the average 0% (expected minimum) and 100% interphase signal (expected maximum) values of three technical replicates. Accuracy of quantification is determined by R^2^ of the observed H3K9ac signal relative to this line of expected signal. **e.** Scatter plots showing H3K9ac signal quantified at 1 kb bins of chromosome 19, in samples with different H3K9ac levels (25%, 50%, 75% and 100% interphase HeLa-S3 cells vs samples with minimal H3K9ac signal (100% mitotic/0% interphase HeLa-S3 cells). ChIP-wrangler-normalized signal is plotted in dark grey, read-depth-normalized signal in light-grey. Additional data in **Supp. Figs. 1-3**, Genome browser session [here](https://genome.ucsc.edu/s/lahodge/H3K9ac_titration_spikenorm_submit).

**Fig. 2.**
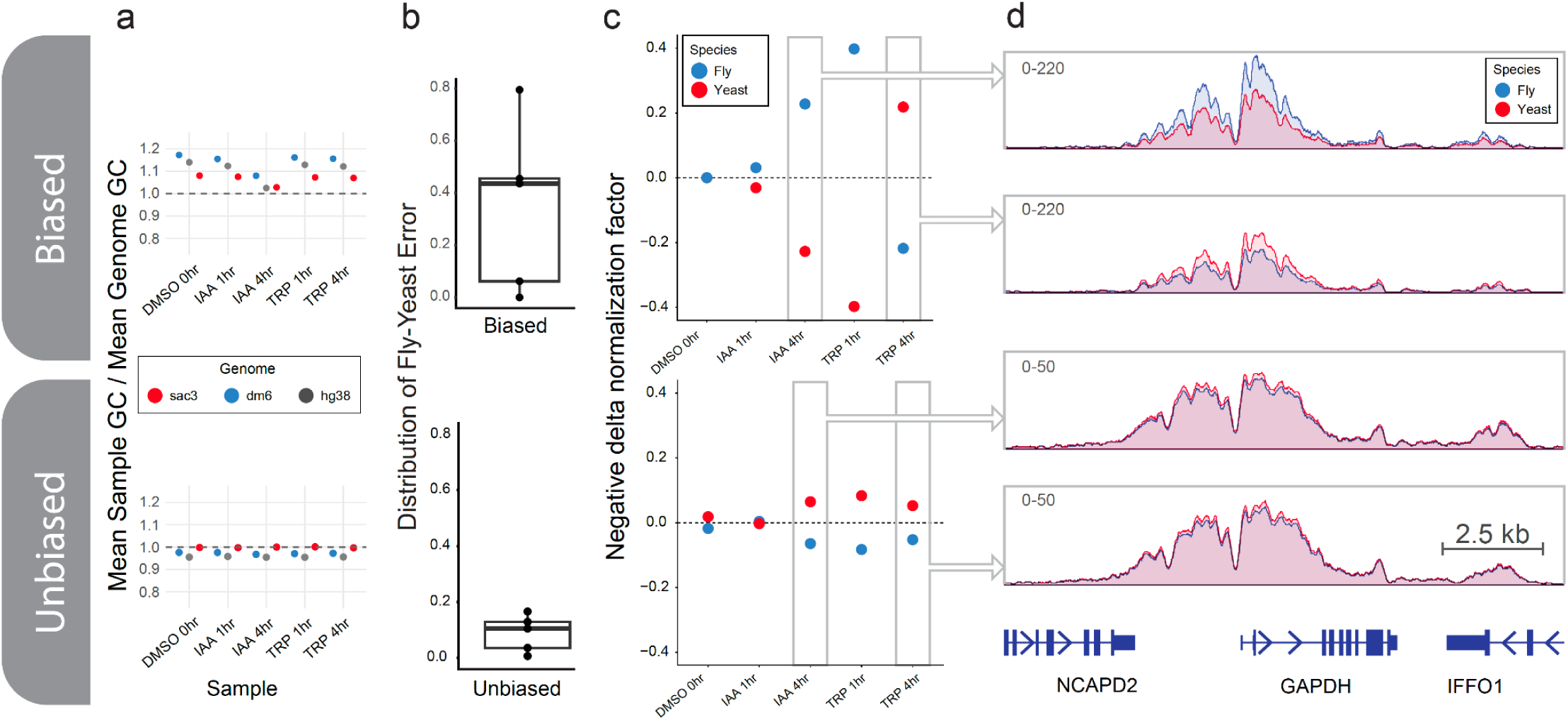
An example of the increased rigor obtained by using spike-ins from two species. **a.** GC content of ChIP-seq input libraries relative to the expected GC content for each species (Blue: *D. melanogaster*, Grey: *H. sapiens*, Red: *S. cerevisiae*). Expected GC contents for each species are 42%, 40.9%, and 38%, respectively. **b.** Boxplots depicting the difference between the average-adjusted normalization factors obtained from each species (**Methods**) for biased (higher than expected genomic GC content) or unbiased libraries from an experiment to test the impact of Triptolide and auxin-degradation of RNAPII on transcription (5 different conditions, see Fig. 4). **c.** The negative delta in spike-in normalization factors compared to expected (expected value: the average of normalization factors for both species), colored by species (blue: fly; red: yeast). The dotted line represents the average normalization factor, depicting the expected agreement between the two spike-in species. **d.** An example genome browser tracks (IGV^7^) showing *H. sapiens* (hg38) H3K27ac signal normalized to either *D. melanogaster* or *S. cerevisiae* (blue: fly; red: yeast). Additional data for this analysis are in **Supp. Figs. 4-5**, genome browser session [here](https://genome.ucsc.edu/cgi-bin/hgTracks?hgS_doOtherUser=submit&hgS_otherUserName=lahodge&hgS_otherUserSessionName=LP88_LP85_spikenorm_speciescompare).

To evaluate the ability of our novel spike-in normalization approach to correctly quantify protein-DNA interactions, we analyzed experimental conditions with a known ground truth. To minimize human error in our benchmarking experiments we employed our fully automated ChIP-seq process^5^. We generated a dataset with predefined levels of histone acetylation to benchmark ChIP-wrangler. We used mitotic cells – where global histone acetylation levels are reduced^6^ – that were mixed with interphase cells at known ratios, where the levels of this mark increase with the proportion of interphase cells. In experiments with good QC metrics (i.e., consistent spike-in quantities between samples), jointly normalizing by the two species provided modest increases in accuracy compared to each species individually (improving from R^2^ = 0.95-0.97 with single species to R^2^ = 0.98 with dual spike-in species; **Supp. Figs. 1-2**), and dramatic increases in accuracy compared to read-depth normalization (R^2^ = 0.07) (**Fig. 1b-e**).

Notably, the key advantage of the dual spike-in approach lies in cases where the two normalization factors do not agree. As a demonstration of how such disagreement enables ChIP-wrangler to provide “guardrails” for failed experiments, we used samples where the library preparation conditions led to a skew in the overall nucleotide composition, with higher than expected GC content. Since the average genomic GC proportion of the *D. melanogaster* and *S. cerevisiae* differs (42% and 38%, respectively), the biased library preparation impacts these differently (**Fig. 2a**), and this in turn leads to a discrepancy in the results of separate normalization to each spike-in. While in well-controlled experiments the two spike-ins yield highly similar results (**Fig. 1b-d**, **Fig. 2 bottom**), the normalization outcomes diverged dramatically in the samples with biased library preparation (**Fig. 2 top, Supp. Figs. 4-5**). Note, this technical artefact would have been difficult to detect with only one spike-in.

To evaluate the robustness of ChIP-wrangler across a range of spike-in conditions, we next used our automated ChIP-seq platform^5^ to conduct an additional titration experiment. We used a mixture of samples to generate libraries with a known ground truth, while simultaneously varying the proportions of spike-in to target chromatin. We varied the amount of *D. melanogaster* and *S. cerevisiae* relative to each other, and the total amount of spike-in chromatin relative to the target chromatin, focusing on four different spike-in/target chromatin ratios (**Fig. 3a**). For each of the four ratios, we also conducted a titration, where we varied the levels of histone H3 lysine 27 acetylation (H3K27ac) across six conditions with an estimated dynamic range of 8.3-fold (**Methods**). We generated this dataset by mixing cells treated with DMSO (baseline H3K27ac) and cells treated for 18 hours with 1 µM TSA (high H3K27ac, **Fig. 3b**). ChIP-wrangler accurately reproduces the standard curve of H3K27ac, with R^2^ values ranging from 0.884 - 0.995. Note, excessive amounts of spike-in chromatin had a negative impact on the quantification accuracy, in a similar manner to our previous observations^4^. In particular, Ratio D – where the spike-in material accounted for ∼20% of the total input chromatin – yielded the lowest R^2^ value (0.884).

**Fig. 3.**
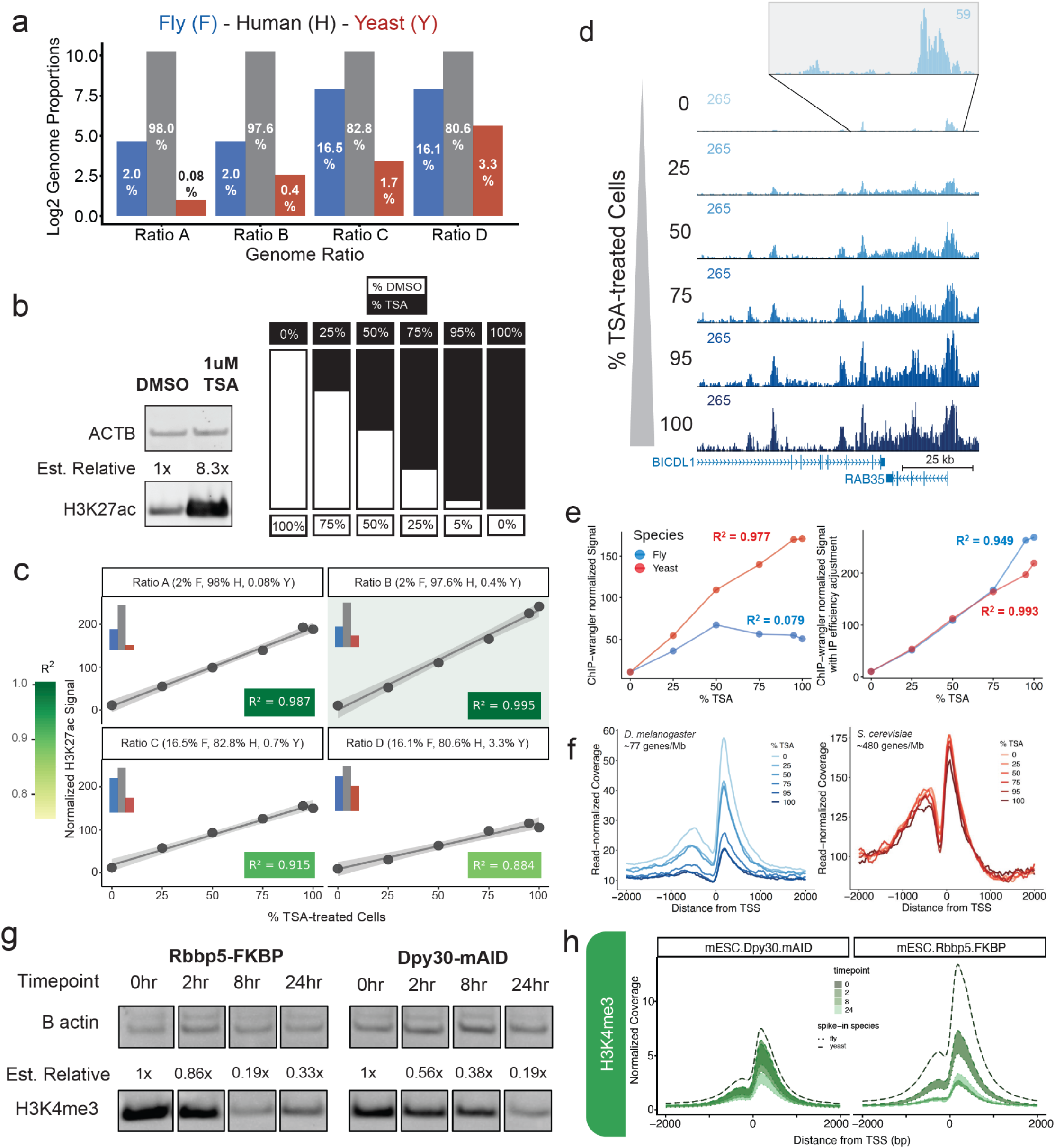
Evaluation of the robustness of ChIP-wrangler across a range of conditions. **a.** Four ratios of target and spike-in chromatin (Ratio A-D) are represented as the log2 of genome proportion (e.g., Ratio A includes 2% Fly (F), 98% Human (H), 0.08% Yeast (Y)). For clarity, we provide both the genomic proportions of spike-ins and target chromatin plotted on the Log2 axis, and the raw expected % of reads for each species in white. **b.** For each of the 4 ratios depicted in (a), H3K27ac signal was titrated across 6 conditions, ranging from baseline (DMSO-treated) to high (1µM TSA-treated) HeLa-S3 cells. The dynamic range of H3K27ac levels was estimated by ImageJ analysis of the Western blot (left), the chromatin was mixed according to the proportions shown by the black and white bars (right). **c.** Titration curves of the 4 ratios described in (a). The ratios are given in the headers and depicted visually using the barplots from (a) for clarity (inserts). For each datapoint in the titration curve, the H3K27ac signal was quantified by the composite signal at all merged peaks (calculated by area under the curve; **Supp. Fig. 6, Methods**). The accuracy of titration curves was defined by R^2^ (bottom right inserts in each plot, the colorbar on the left provides the shades according to R^2^ of each plot). Ratio B (green background) provided the highest accuracy. **d.** ChIP-wrangler normalized UCSC genome browser tracks of the best-performing ratio (Ratio B), showing an increase of H3K27ac along titration conditions. QCs and additional data are in **Supp. Figs. 6-9**, genome browser tracks [here](https://genome.ucsc.edu/cgi-bin/hgTracks?hgS_doOtherUser=submit&hgS_otherUserName=lahodge&hgS_otherUserSessionName=251116_TSAtitration_Hela_Rpb1). A view of the baseline H3K27ac levels with a reduced scale (59 vs 265) is presented on top. **e.** Titration of H3K27ac from Ratio B was normalized using the calculation of a spike-in normalization factor (IP/input), left, or normalized after accounting for IP efficiency estimated using spike-in ChIP-seq signals (right, details in **Methods**). Accuracy of normalization to each species (*D. melanogaster*, blue; *S. cerevisiae*, red) was determined by calculating R^2^ of each resulting curve (plot inserts). **f.** Histogram of H3K27ac at *D. melanogaster* (left) and *S. cerevisiae* (right) promoters, colored by the levels of H. sapiens H3K27ac, varied by mixing TSA-treated (high H3K27ac) and DMSO-treated (low H3K27ac) in different ratios. Note the sensitivity of the signal for *D. melanogaster*, a species with lower gene density compared to *S. cerevisiae* which is mostly unaffected. **g.** Immunoblots showing levels of H3K4me3 at 0, 2, 8, and 24 hr of induced degradation of Rbbp5 (mESC-Rbbp5-FKBP) and Dpy30 (mESC-Dpy30-mAID) lines. β-actin (ACTB) was used as the loading control for each condition. **h.** Histograms of ChIP-wrangler normalized coverage of H3K4me3 signal at ± 2 kb of TSS for 0, 2, 8, and 24 hr of induced degradation of mESC-Dpy30-mAID (left) and mESC-Rbbp5-FKBP (right) lines. Data were normalized to *D. melanogaster* (dotted lines) and *S. cerevisiae* (dashed lines). Note, results from each spike-in species were normalized relative to their respective control condition (0hr) therefore these samples lack a confidence interval estimate (details in **Methods**). QCs and additional data in **Supp. Figs. 6 & 9**.

When normalizing the H3K27ac titration dataset to either spike-in species, we observed that while *S. cerevisiae* provided high accuracy (R^2^ = 0.98), normalization to *D. melanogaster* appeared problematic (R^2^ = 0.08; **Fig. 3e, left**). To correct this issue we estimated the relative IP efficiency for each spike-in species, and adjusted the normalization factors accordingly. This correction improved the agreement between the spike-ins and the overall accuracy of normalization (*D. melanogaster* R^2^ = 0.95, *S. cerevisiae* R^2^ = 0.99, **Fig. 3e, right, Methods**). In this case, the H3K27ac IP efficiency was minimally impacted in *S. cerevisiae* but showed a strong decrease in *D. melanogaster* with an increase in H3K27ac epitope levels, highlighting the differences in the properties of the spike-in chromatin from these two species (**Fig. 3f**, **Discussion**).

We additionally evaluated ChIP-wrangler’s ability to detect a global change in ChIP-seq signal using mouse (*M. musculus*) instead of human samples as the target species. We employed mouse embryonic stem cells (mESCs) that were genetically modified to allow inducible degradation of two of the COMPASS complex members – Rbbp5 (mESC-Rbbp5-FKBP) and Dpy30 (mESC-Dpy30-mAID). These lines were previously used to demonstrate the reduction in H3K4me3 levels following depletion of these COMPASS subunits^8^. We induced the degradation of Rbbp5 and Dpy30 (either using dTag or auxin, respectively) for 0, 2, 8, and 24 hours and applied ChIP-wrangler using *D. melanogaster* and *S. cerevisiae* spike-ins. Quantification of H3K4me3 signal by Western Blot identified a reduction in H3K4me3 levels (3-fold and 4.2-fold reduction after 24 hours of Dpy30 and Rbbp5 depletion, respectively) following the degradation of the COMPASS subunits (**Fig. 3g**). We observed a similar degree of reduction in H3K4me3 levels in the normalized ChIP-seq data (**Fig. 3h**) demonstrating the applicability of our dual spike-in approach to mouse cells along with its ability to detect transitional changes.

Two recent studies have challenged the model that histone acetylation is a regulatory mechanism that occurs upstream of transcription^9^, positing instead that histone acetylation is dependent on RNAPII activity^10,11^. These conclusions were based on using small-molecule inhibition of RNAPII transcription, which led to the appearance of a drastic decrease in global histone acetylation levels. Previously, we and others noted that the genomic conclusions of these studies have potentially been affected by experimental artefacts such as low and variable spike-in chromatin amounts^4,12,13^. This is further supported by a recent study that showed that global histone acetylation does not decrease upon RNAPII inhibition by quantitative mass spectrometry^14^. To address the discrepancy between these studies, and as a proof of concept for our new method, we tested the impact of acutely inhibiting transcription on a panel of histone modifications. We used two complementary methods to inhibit transcription by RNAPII for 1 or 4 hours in HCT116 cells. First, as in Wang et al.^10^, we treated cells with Triptolide, which blocks TFIIH recruitment leading to inhibition of transcription and the degradation of RNAPII^15^. In parallel, we established an HCT116 cell line with an auxin-inducible degron fused to the RPB1 N-terminus to deplete RNAPII upon auxin treatment (**Fig. 4a**).

**Fig. 4.**
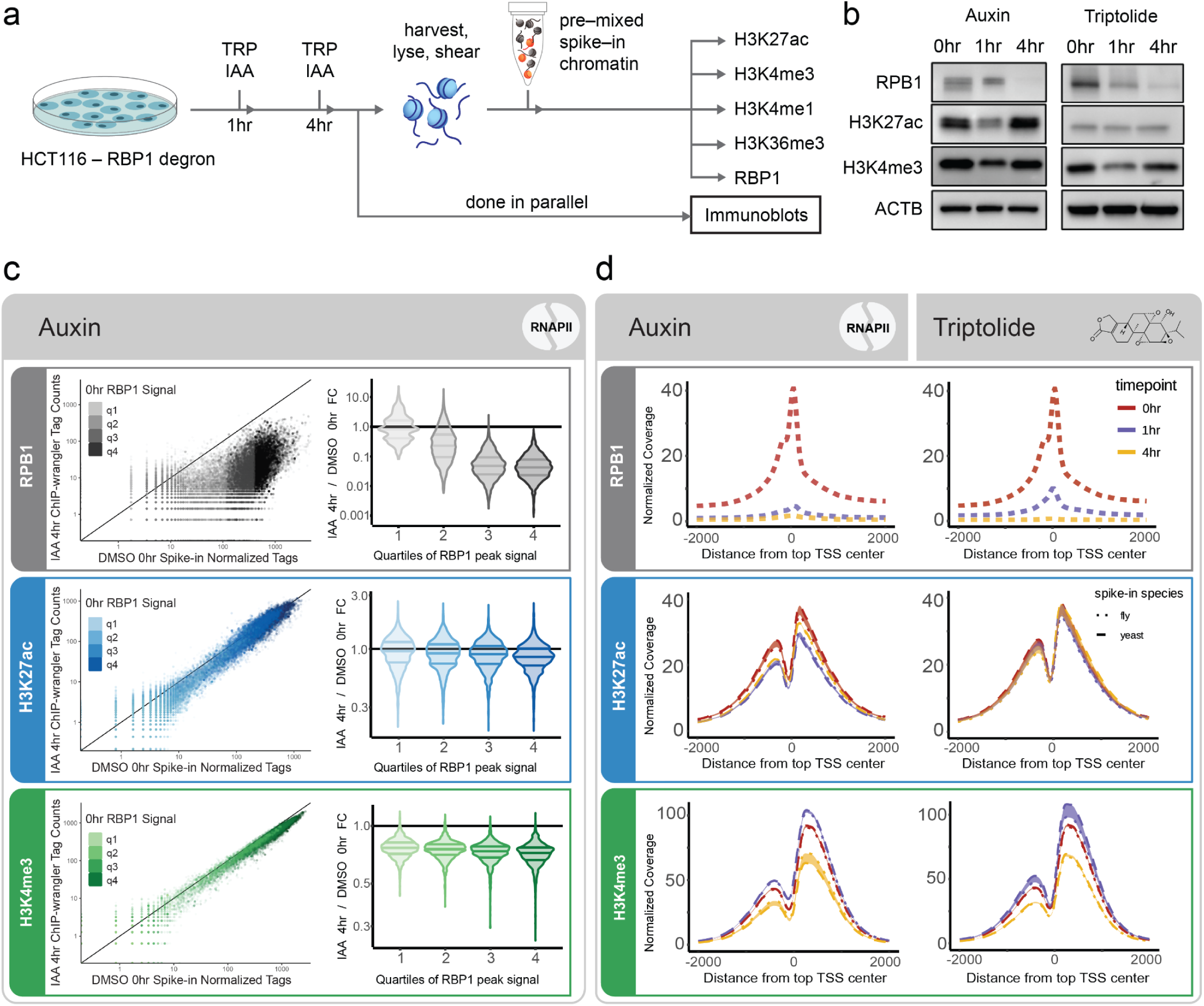
Acute depletion of RNAPII shows a modest impact on promoter-proximal active histone modifications. **a.** Overview of the experimental design for depletion of RNAPII in HCT116 cells by using two orthogonal approaches: auxin-induced degradation of RPB1 (IAA) or a small molecule inhibitor (Triptolide, TRP) that leads to degradation of RNAPII. **b.** Immunoblots showing levels of RPB1 (the catalytic subunit of RNAPII), H3K4me3, H3K27ac and loading control (ACTB) at each time point. **c.** Comparison of ChIP-wrangler normalized RPB1, H3K27ac, and H3K4me3 signal at promoters (defined as TSS +/- 2kb) after 4 hours of auxin treatment relative to untreated levels, shaded according to the quartiles of untreated RPB1 signal (q1-q4, inserts), displayed as violin plots on the right. Note: RPB1 data were normalized to *D. melanogaster* only (**Supp. Fig. 8**, **Methods**). **d.** Histograms of ChIP-wrangler normalized RPB1, H3K27ac, and H3K4me3 signal at active promoters (top 50% of RPB1 signal) at 0, 1, and 4 hrs of treatment by auxin and Triptolide (header labels). Data were normalized to *D. melanogaster* (dotted lines) and *S. cerevisiae* (dashed lines). Note: RPB1 data were normalized to *D. melanogaster* only (**Supp. Fig. 8**, Methods). Additional data is presented in Supplemental Figs. 10-13, and the genome browser session for HCT116 data is [here](https://genome.ucsc.edu/cgi-bin/hgTracks?hgS_doOtherUser=submit&hgS_otherUserName=lahodge&hgS_otherUserSessionName=LP88_HCTrep1_spikenorm).

Both Triptolide treatment and auxin-induced degradation significantly decreased RNAPII protein levels in whole cell lysates (measured by immunoblots) and bound to chromatin (measured by ChIP-seq) after 1 hour, with a pronounced reduction after 4 hours (**Fig. 4b-d, Supp. Figs. 10-12**). Immunoblots for total H3K27ac and H3K4me3 show a transient reduction in signal after 1 hour of RNAPII depletion, which increases back to almost pre-treatment levels after 4 hours (**Fig. 4b**). Further, immunoblots for the enhancer mark H3K4me1 and the gene body transcription associated H3K36me3 show that their global signal is mostly stable after 1 and 4 hours of RNAPII depletion (**Supp. Material**).

Next, using ChIP-wrangler we analyzed the changes in genomic patterns of each histone modification. As opposed to the previous observations showing a drastic reduction in promoter H3K27ac^10,11^ and H3K4me3^11^ following small molecule treatment, the overall levels of H3K27ac at transcription start sites (TSSs) were mostly maintained following the depletion of RNAPII (**Fig. 4c-d**). Further, promoter levels of H3K4me3 showed a small but notable increase after 1 hour of RNAPII depletion, followed by a slight reduction in signal after 4 hours of treatment, resembling trends observed in a recent report^16^. In addition, H3K4me1 levels remained relatively constant, while H3K36me3 slightly decreased along gene bodies in a gene-length dependent manner after 4 hours of treatment by either Triptolide or auxin-mediated degradation of RNAPII (**Supp. Figs. 10-12**).

While the general levels of histone marks associated with regulatory regions were relatively static (**Fig. 4c-d, Supp. Figs 7-9**), it is also apparent that a subset of genomic regions exhibited changes following transcriptional inhibition. In addition, changes in H3K27ac after HDAC inhibition with TSA revealed striking increases of H3K27ac along active gene bodies (**Fig. 3d**, **Fig. 5a-b**) and genes that lack RNAPII binding (measured by RPB1), show no increase in H3K27ac (e.g. *NLRP9*, **Fig. 5b** right). Together these observations indicate a relationship between active H3K27ac deposition and transcription elongation. Overall, H3K27ac changes were highly correlated between both RNAPII inhibition approaches that we used (**Fig. 5c**). Further, among the top 50% of H3K27ac peaks, those increasing following perturbation of RNAPII were enriched for promoter-proximal regions, while loci that show a decrease in H3K27ac were mostly composed of promoter-distal peaks (**Fig. 5c**). Interestingly, promoter-distal peaks found within gene bodies show a lower degree of H3K27ac reduction compared to those in intergenic regions (**Fig. 5d,** p = 2e-16, Wilcoxon rank sum test, two-sided). Pairwise comparison of changes in H3K27ac signal in all genomic regions were statistically significantly different, with one exception (**Supp. Fig. 15**). Altogether, this suggests that the presence of elongating RNAPII could potentially impact the maintenance of acetylation levels in transcribed regions, in line with previous observations showing the association of the histone acetyl-transferases PCAF with transcribing RNAPII^17^.

**Fig. 5.**
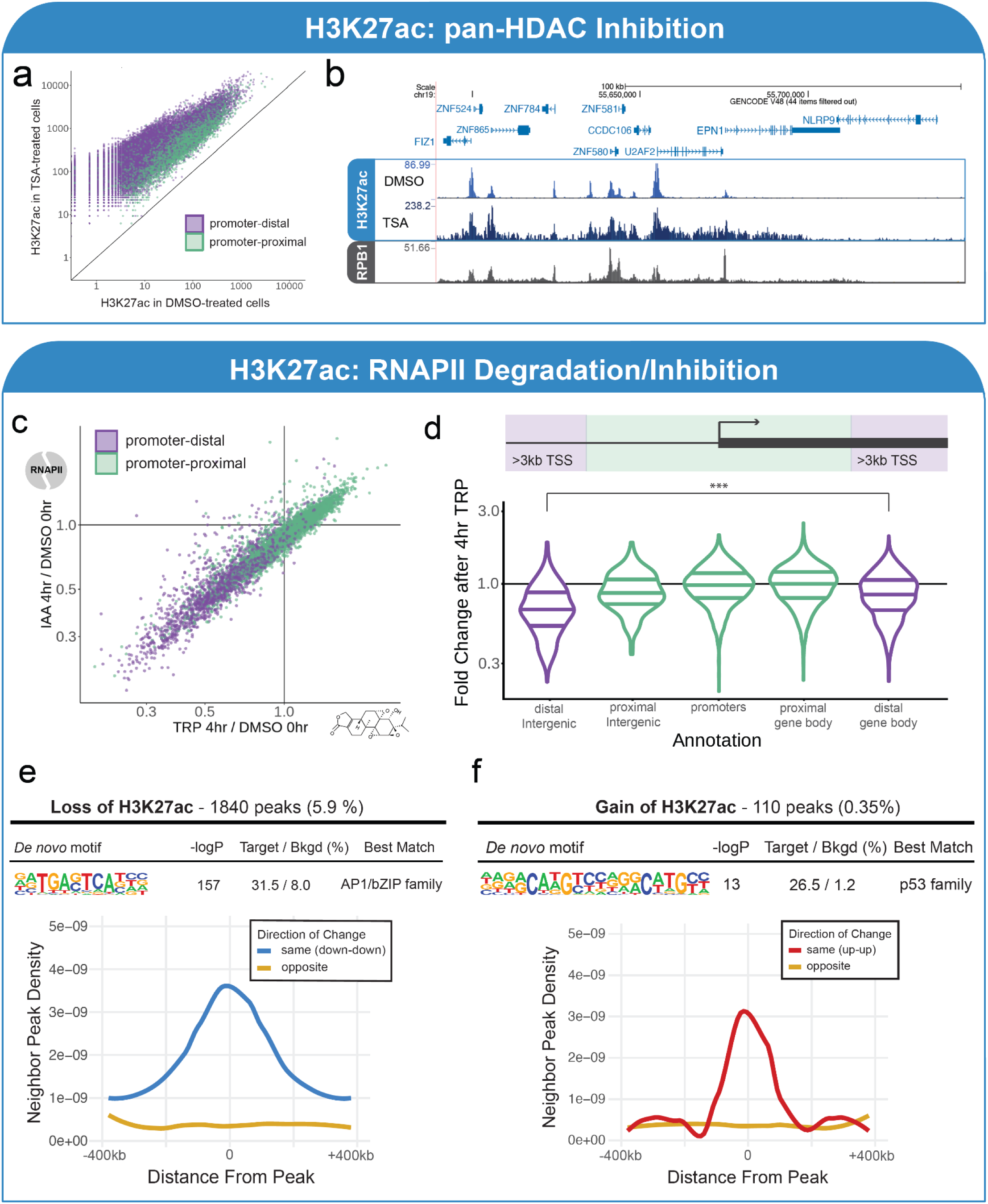
Changes in H3K27ac after RNAPII degradation are modest, yet positionally-specific. **a.** Promoter-distal peaks (defined as >3kb from TSS, purple) show a greater increase in H3K27ac following treatment of HeLa-S3 cells with pan-HDAC inhibitor (TSA) compared to promoter-proximal (defined as <3kb from a TSS, green). **b.** UCSC genome browser tracks of ChIP-wrangler normalized H3K27ac signal from HeLa-S3 cells treated with TSA or DMSO (top); and ChIP-seq for RBP1 of the same cells (bottom). Note the difference in y-axis for DMSO and TSA-treated H3K27ac tracks due to the major differences in the signal between these conditions. **c.** Scatterplot showing correlation of change in H3K27ac after 4 hours of auxin and Triptolide at peaks in the top 50% of untreated signal, colored by distance of peaks to TSS as in (a). **d.** Of the promoter-distal peaks in (c), peaks not on gene bodies show a greater decrease in H3K27ac signal than genic peaks (p = 2e-16, Wilcoxon rank sum test, two-sided). Here we highlight the difference between distal Intergenic and distal gene body peaks, all except one of the permutations between all genome annotations were statistically significant, and are provided in **Supp. Fig 15**. **e-f.** Differential H3K27ac peaks as determined by DESeq2 after 4 hours of both Triptolide and auxin treatment, only peaks differentially changing >2 fold for both treatments were kept. Motifs downregulated after TRP/IAA mainly included a general enhancer repertoire, while upregulation of the p53 motif family indicates activation of cellular stress response. **e.** H3K27ac peaks decreasing after RNAPII inhibition or degradation (1840, 5.9% of peaks) are near other decreasing peaks. **f.** H3K27ac peaks increasing after RNAPII inhibition or degradation (110, 0.35% of peaks) are near other increasing peaks. Data for **a-b.** are ChIP-wrangler normalized H3K27ac and Rpb1 in HeLa-S3 cells; additional data is in **Supp. Fig. 14**, genome browser tracks [here](https://genome.ucsc.edu/cgi-bin/hgTracks?hgS_doOtherUser=submit&hgS_otherUserName=lahodge&hgS_otherUserSessionName=251116_TSAtitration_Hela_Rpb1). Data in **c-f.** are ChIP-wrangler normalized H3K27ac in Triptolide and auxin-treated HCT116-RPB1 degron; additional data is in **Supp. Fig. 16-17**, genome browser tracks [here](https://genome.ucsc.edu/cgi-bin/hgTracks?hgS_doOtherUser=submit&hgS_otherUserName=lahodge&hgS_otherUserSessionName=LP88_HCTrep1_spikenorm).

To further characterize the regions impacted following perturbation of transcription, we identified significantly differentially regulated H3K27ac peaks using DESeq2^18,19^ with ChIP-wrangler-derived size factors (**Methods**). We first conducted DESeq2 analysis (Fold Change > 2, adjusted p-value < 0.05) for each of the auxin and Triptolide experiments. Then, to study the regions that are most confidently impacted by loss of RNAPII, and to exclude regions that are subject to perturbation-specific effects, we restricted our analysis to only the set of overlapping regions. Out of the merged set of 31,075 H3K27ac peaks obtained from the 4-hr timepoints of both perturbations and controls, 1840 (5.9%) had reduced signal, and 110 (0.35%) had increased signal (**Fig. 5**, Supp. Fig. 16).

Analysis of DNA motifs revealed a strong enrichment for the transcription factor family AP-1/bZIP in peaks that show a reduced H3K27ac signal (**Fig. 5e**). Members of the AP-1/bZIP transcription factor family are known to be associated with cis-regulatory elements (enhancers)^20–22^. The enrichment of this DNA motif family in the set of peaks with reduced H3K27ac aligns with our observation that such promoter-distal elements show a reduced signal following transcription perturbation (**Fig. 5e**), which make up 82% of DESeq-identified peaks decreasing in both treatment conditions. In contrast, peaks gaining acetylation were enriched for p53 response elements (**Fig. 5f**). This enrichment is potentially linked to cellular stress pathways activated by loss of RNAPII^23^, and is consistent with previous observations that activating transcription factors can increase acetylation near their binding sites in the absence of RNAPII^16^. Lastly we observed that regions of differential acetylation that are adjacent to one another appear to be coordinately regulated within genomic neighborhoods; peaks which show a decrease in signal following perturbation of transcription are found closer to other peaks that also show a reduction in histone acetylation, and vice versa (**Fig. 5e-f** bottom).

## Discussion

We presented here ChIP-wrangler, which jointly incorporates molecular and computational approaches to enable rigorous normalization of ChIP-seq datasets and quantitative detection of variations between conditions. We first benchmarked our normalization approach by using a combination of titration and mixing experiments that allowed us to generate datasets with known ground truth. We then used ChIP-wrangler to reevaluate previous claims that active histone modifications are dependent on transcription. We observed that perturbation of RNAPII had a minor impact on the levels of H3K4me3 and H3K27ac, in contrast to previous claims^10,11^. For instance, only ∼6% of the H3K27ac peaks showed a consistent reduction across transcription perturbation systems, of which promoter-proximal and -distal regions are differentially impacted by transcriptional inhibition.

While earlier versions of spike-in normalization approaches enable sensitive detection of global changes in signal, there are potential risks in implementing these methods. Similar to most molecular biology protocols, the generation of spike-in normalized ChIP-seq libraries include several weak points where technical biases/artefacts can be introduced without the user’s awareness^4^. Such weak points include the possible error inherent in introduction of exogenous chromatin (e.g., variation between the amount spiked to each sample), or artefacts introduced during library preparation such as GC bias or read length variation that skews the resulting data.

As the normalization process relies on the ratio of reads originating from the spike-in and target chromatin, proper quantification of the scaling factor derived from the ratio is critical for processing of the data. Thus, in a similar manner to the increased level of robustness achieved by redundancy of key components in other fields such as engineering and computer science, ChIP-wrangler uses independent exogenous chromatin preparations from two different species. In the current implementation we used a pre-mixed preparation of chromatin from *D. melanogaster* and *S. cerevisiae*, which have different genomic properties (e.g., genome size, chromatin organization, average GC content, etc). The additional spike-in serves as a failsafe as it provides means for additional QC steps that are not available when using a single spike-in (e.g., **Fig. 2**, further discussed below).

Additionally, ChIP-wrangler estimates the efficiency of the immunoprecipitation (IP efficiency), which can be a major source of variation between ChIP-seq samples in an experiment (e.g., **Fig. 3e,f**). The deviation in IP efficiency can vary due to several experimental factors. These include variation in non-specific antibody binding, or large discrepancies in the amount of epitope, leading to antibody competition^24^ (**Supp. Fig 18, GitHub readme**). Here, we noticed that when we used human samples with high H3K27ac levels (following pan-HDACs inhibition), the signal-to-noise ratio in *D. melanogaster* spike-in was strongly impacted by the levels of H3K27ac in the target (H. sapiens). On the other hand, the *S. cerevisiae* spike-in was mostly insensitive to changes in human epitope abundance (**Fig. 3f**). We hypothesize that this variation between the two species is in some part due to the underlying differences in gene density between the two organisms (77 vs. 480 genes/Mb in *D. melanogaster* and *S. cerevisiae*, respectively). This aligns with our recent study^5^, where we observed that weak ChIP-seq signals show higher sensitivity to the ratio of antibody to epitope, while loci with strong signal are less sensitive, and are detected in a similar manner across a wide range of antibody to epitope ratios.

We quantified the accuracy of ChIP-wrangler via two comprehensive cell mixing/titration experiments, either increasing or decreasing the levels of two histone acetylations (H3K9ac and H3K27ac) by separate systems. To reduce human error in the high number of samples required for our benchmarking experiments we employed our fully automated ChIP-seq process^5^. Our analyses showed that ChIP-wrangler accurately captures the signal across a range of conditions. These titration/mixing experiments also allowed us to define an optimal ratio of *D. melanogaster*, *S. cerevisiae* and the *H. sapiens* target chromatin (**Fig. 3**).

We demonstrated the importance of the dual spike-in approach and the value provided by including genomes with different average GC content (*D. melanogaster*, 42%; *S. cerevisiae*, 38%). When analyzing ChIP-seq libraries with a biased nucleotide composition (technically flawed), we obtained opposing biological conclusions depending on the spike-in that was used for normalization. Thus, if we used the *D. melanogaster* spike-in, histone acetylation appeared to decrease after 4 hours of transcription inhibition, whereas if we normalized to *S. cerevisiae* the acetylation signal increased (**Fig. 2**). Including spike-ins from both species immediately alerted us to the discrepancy in our results.

Resolving the issue of whether histone acetylation is a consequence of RNAPII activity or contributes to transcription regulation is important for interpreting histone modification data and future investigations of regulatory mechanisms. Using ChIP-wrangler, we have demonstrated that RNAPII activity is not necessary for the maintenance of these histone modifications, and that the relationship between transcription and active histone marks is nuanced and coordinated by multiple mechanisms.

Although these changes were only observed in a small subset of H3K27ac peaks (∼6%), focusing on these regions we observe a differential response to the loss of transcription. Only 110 regions showed an increase in acetylation, and these were enriched for gene promoters related to cellular stress-response (**Fig 5**). Of the regions with decreasing H3K27ac (5.9% of all peaks) most were putative enhancers, and were enriched for the presence of AP-1 (bZIP) DNA motifs. On the other hand, promoter-distal peaks showed the highest increase in H3K27ac after inhibition of HDACs, particularly along gene bodies (**Fig. 5**). This suggests that promoter-distal peaks are more dependent on ongoing transcription and the presence of elongating RNAPII than promoter-proximal peaks (**Fig. 5f**) in a similar manner to recent findings^16^. Our previous observations of the colocalization of HATs, HDACs and RNAPII^25^ further support this model of coordinated regulation. Together, this suggests that chromatin context and transcriptional activity jointly influence H3K27ac regulation. Notably, our study only addresses the role of transcription in the maintenance of histone marks at limited time points, and does not investigate the establishment or the dynamics of these histone modifications.

In our study we used ChIP-wrangler to quantify the changes in histone modifications using two inducible degradation systems to perturb chromatin regulators, namely the auxin inducible degron (AID) system to target RPB1 and Dpy30 and the dTag system to enable inducible degradation of Rbbp5. As experiments using these and similar systems for rapid degradation of regulators to establish causality are becoming common^26,27^, ChIP-wrangler could enable accurate quantification of either global or more selective histone modification changes following the depletion of the targeted protein.

We detail below considerations to increase robustness when applying ChIP-wrangler: (***i***) whenever possible, all experimental perturbations should be conducted in parallel, and downstream library preparation of multiple biological replicates should be done simultaneously to minimize batch effects. (***ii***) Paired-end sequencing enables estimations of fragment length and improves genomic alignment between species. (***iii***) The IP efficiency defined above, is a critical QC for evaluating the success of the ChIP-seq for the spike-in samples as well as for calibration of the normalization factors. Thus, the species used for spike-in should be chosen such that the antibody recognizes all target epitopes. (***iv***) When using spike-in from species with genomes that are closely related to the target species (e.g., mammalian spike-ins for a human target), sequencing with longer read lengths and paired end sequencing could reduce the chances of ambiguously aligned reads.

One of the limitations of ChIP-wrangler is that in the current implementation, using *D. melanogaster* and *S. cerevisiae,* is mostly amenable to conserved epitopes such as the histone modifications H3K27ac and H3K4me3. Notably, it is possible to use spike-ins that are closely related to the target species. Another limitation is that to avoid dubious alignments/multimappers and reduce the chances of misquantification, all multimapping reads are discarded. This can potentially reduce the total reads aligned to each species.

Lastly, while ChIP-wrangler was developed for normalization of ChIP-seq data, in theory, this concept can be applied to other next generation sequencing (NGS) approaches where incorporation of a failsafe step can allow increased robustness.

## Supporting information

patel-etal-biorxiv-sup-figs-revision2-submit

patel-etal-biorxiv-sup-material-revision2-submit

Patel-etal-biorxiv-Supplemental-Table-revision2 -submit

## Acknowledgements

Research reported in this publication was supported in part by NIH/NIMH grants R01MH127077 (A.G. and C.B.), R35GM149520 (C.B.), R01GM129523 (S.H.), NSF grant 2003358 (A.G.) and BSF grant 2019688 (I.S.). T.X. and Y.C. were supported in part by the CIRM Research Training Grant EDUC4-12804. Ed.M was supported in part by an institutional award to the UCSD Genetics Training Program from the National Institute for General Medical Sciences, T32 GM145427. We thank Kristian Helin and Zheng Fan for their technical assistance and use of their mESC-Dpy30-mAID and mESC-Rbbp5-FKBP cell lines. We thank R. Wachs for her help with the illustrations, Y. Farjoun for his scientific input and Camila De Arruda Saldanha for technical assistance creating the Rbp1-auxin-inducible degron cell line. We thank the Pillus lab at UC San Diego for providing *S. cerevisiae* cells and guidance on yeast culture and ChIP-seq and the Saphire lab at La Jolla Institute for Immunology and Joiner lab for providing *D. melanogaster* S2 cells. We thank the Stem Cell Genomics Core at the Sanford Stem Cell Institute and the Epigenomics Core at UC San Diego for providing sequencing services.

## Methods

### HCT116 Cell Media and Culture

HCT116-EF1a-OsTIR1-Rpb1 cells were cultured in McCoy’s 5A media (Thermo Scientific 16600082) with 10% head-inactivated fetal bovine serum (Life Technologies 10438026) and 1x antibiotic-antimycotic. Cells were incubated at 37°C and 5% CO2.

### HeLa-S3 Cell Media and Culture

HeLa-S3 cells were grown in Dulbecco’s Modified Eagle’s Medium (DMEM) supplemented with 10% head-inactivated fetal bovine serum (Life Technologies 10438026), 1x antibiotic-antimycotic (Thermo Scientific 15240062), L-glutamine (Invitrogen A2916801), and 110mg/L sodium pyruvate (Gibco 11360070). Cells were incubated at 37°C and 5% CO_2_.

### mESC Cell Media and Culture

mESC-Dpy30-mAID and mESC-Rbbp5-FKBP cell lines were obtained from the Helin lab. Cells were cultured on 0.2% gelatin (Sigma-Adlrich G1393) plates, precoated for at least 2 hours at 37C. mESCs were grown in Glasgow Minimal Essential Medium (GMEM) supplemented with 15% fetal bovine serum (FBS), 10uM sodium pyruvate, 1x MEM-NEAA (Gibco 11140-050), 50uM β-mercaptoethanol (Thermo Fisher Scientific 31350010) and 1x pen-strep (Thermo Fisher Scientific 15-140-122).

### S2 Cell Culture

Drosophila S2 cells were grown in Schneider’s Drosophila Medium (Gibco 21720024) + 10% FBS at 28°C in the dark.

### Yeast Cell Culture

From frozen stocks, starter cultures of strain LPY6494 were inoculated in 3mL of liquid YPD media (Gibco A1374501), grown to log phase at 30°C at 220rpm, then expanded to 50mL cultures at an OD of 0.08 and grown for 16 hours at 30°C and 220rpm. OD was estimated with NanoDrop UV-Vis Spectrophotometer (Thermo Scientific 13-400-519). Yeast culture and fixation performed as previously described^28^.

### Hela-S3 Mitotic Synchronization

Cells were synchronized in G1/S by adding 2 mM thymidine (Sigma T1895-1G) for 16 hr, washed with PBS, released for 3 hr in fresh medium, and arrested in pro-metaphase using 10 µM S-trityl-L-cysteine (STC) (Sigma 164739) for 12 hr then harvested.

### Treatments

Cells were treated with 500nM Triptolide (Sigma-Aldrich T3652), 500uM 3-indole-acetic acid/IAA (Sigma Aldrich I3750) or 1uM TSA (Selleck Chem S1045). All small molecules were reconstituted in DMSO.

### Immunoblotting

Cells were lysed in RIPA lysis buffer for 30 minutes on ice, vortexing every 10 minutes. Lysates were sonicated on PIXUL (Active Motif 53130) for 24 minutes, then clarified by centrifuging at 14000g for 10 minutes, at 4°C. Protein concentration was determined using a DC protein assay (Bio-Rad 500-0116). Samples were diluted in Protein Sample Buffer. Lysates were run on a NuPAGE 4-12% Bis-Tris Gel (Invitrogen NP0321), running at 200V for 30 minutes in 1x NuPAGE MES SDS running buffer (Invitrogen NP0002). Protein was transferred to 0.45μm PVDF membrane (Millipore IPVH00010) at 100V for 90 minutes in 1x NuPAGE Transfer Buffer (Invitrogen NP00061) with 20% MeOH (Carolina Biological Supply 874913). Ponceau staining (Sigma-Aldrich P7170) was performed to confirm transfer of protein, then the membrane was washed three times with TBST buffer. The membrane was incubated in Blocking Buffer for 1 hour, then washed twice with TBS, and once with TBST, each for 5 minutes. Primary antibodies are diluted according to manufacturer’s recommendations in Antibody Dilution Buffer, then incubated overnight at 4°C on a benchtop shaker. Washes were done for 5 minutes, twice in TBS, then once in TBST, then membranes were incubated for 1 hour in secondary antibodies, green or red. Membranes were finally washed twice in TBS, once in TBST, and imaged with the Bio-Rad ChemiDoc Imaging System.

### Buffer components

- RIPA Lysis Buffer: 1x RIPA buffer (diluted from 10x RIPA Buffer, Cell Signaling Technologies 9806), 1x Protease Inhibitor (Roche 11873580001), 1x PMSF (Cell Signaling Technologies 8553), 1x phosphatase inhibitor (Roche 04906845001)
- Protein Sample Buffer: 1x NuPAGE LDS Sample buffer (Invitrogen NP0007) and 4% β-mercaptoethanol (Sigma-Aldrich M3148)
- TBST: 8.8 g NaCl, 2.4 g Tris HCl, 0.56 g Tris Base, 500uL Tween in 1L
- Blocking Buffer: PBS (Gibco 10010-023) + 5% BSA (Bovine Serum Albumin Fraction V, Fisher Bio BP1600-100)
- Antibody Dilution Buffer: PBS (Gibco 10010-023) + 5% BSA (Bovine Serum Albumin Fraction V, Fisher Bio BP1600-100) + 2.5% Sodium Azide (VWR UN3287).

### Antibodies

- CST H3K27ac (D5E4): 1:1000
- CST H3K4me3 (C42D8): 1:1000
- CST H3K4me1 (D1A9): 1:1000
- CST H3K36me3 (D5A7): 1:1000
- CST Rpb1 NTD (D8L4Y): 1:1000
- abcam B-actin (8226): 1:850
- 2’ green 680RD LI-COR, IRDye 680RD Donkey anti-Mouse cat # 926-68072
- 2’ red 800Cw IRDye 800CW Goat anti-Rabbit cat # 926-32211

### ChIP-seq

ChIP-seq was performed as previously described^29^, all buffer components listed below. To allow for processing of many samples in parallel, we use our automated ChIP-seq platform^5^, however for clarity we describe manual steps below. Briefly, HCT116 or HeLa-S3 cells were double crosslinked on plates with 2 mM DSG for 30 minutes (Fisher Scientific NC1736108) and 1% formaldehyde (Pierce ThermoFisher PI28906) for 10 minutes. Crosslinking was quenched with 1/20th volume of 2.625 M glycine (EMD Millipore 357002) and 0.5% BSA (NEB B9000S). Cells were washed by resuspending 3 times with 0.5% BSA/PBS + Protease inhibitor (PIC, Roche 11697498001), spinning down at 1200g for 5 minutes at 4°C and removing the supernatant. Washed pellets were snap frozen in dry ice then stored in -80°C. Cells were lysed with LB3 lysis buffer + PIC, then chromatin was sheared on the PIXUL sonicator (Active Motif 53130) for 108 minutes. S2 cells were crosslinked in 1% formaldehyde for 10 minutes at room temperature, then quenched with glycine and 10% BSA to a final concentration of 0.5%, lysed in LB3 buffer, and sonicated on the PIXUL for 60 minutes. *S. cerevisiae* cultures in log phase were crosslinked with 0.86% formaldehyde for 30 minutes on a room temperature rotating shaker. The cell wall was digested with Zymolase at 37°C for 60 minutes, cells were washed by pelleting 3x at 5000g and washing in PBS + Protease inhibitor, then resuspended in NP-S buffer + PI. Sheared *S. cerevisiae* chromatin was sonicated on the PIXUL for 108 minutes. All PIXUL sonication was done with settings Pulse (N) = 50, PRF (kHz) = 1.00, Burst Rate (Hz) = 20, only Process Time was varied. Sheared chromatin from each species were clarified by centrifugation at 15000g, for 10 minutes at 4°C, then combined and inputs were taken before immunoprecipitation for each condition, to assess the relative ratios of chromatin. To HeLa-S3 cells, approximately 100k S2 and 500k *S. cerevisiae* cells were added per immunoprecipitation reaction, unless otherwise specified (**Fig. 3**). To HCT116 cells, approximately 500k S2 and 100k *S. cerevisiae* cells were added per 1M cells in each immunoprecipitation reaction.

To each IP, 10 µL of Protein A beads (Thermo Fisher Scientific 10002D), 1% Triton X-100 and 1x PIC were added. 1 million HCT116 cells or HeLa-S3 cells were used per immunoprecipitation reaction. Below are the volumes per IP of each antibody used:

- CST H3K27ac (D5E4): 1 µL
- CST H3K4me3 (C42D8): 1 µL
- CST H3K4me1 (D1A9): 1 µL
- CST H3K36me3 (D5A7): 1 µL
- CST Rpb1 NTD (D8L4Y): 4 µL
- CST H3K9ac (C5B11): 4 µL

Samples were incubated overnight on a rotator at 4°C.

IP Washes were performed: 3x with 180 µL of WBI, 3x with 180 µL of WBIII, and 2x with 185 µL TET, then eluted in 25 µL of cold TT Buffer. Input chromatin containing target and spike-in species’ chromatin from each biological condition were diluted 1:62.5x in TT buffer to a final volume of 25 µL. Libraries were prepared with the NEB Ultra II Library Prep Kit (NEB E7645L). NEXTflex Dual-Barcode ChIP-seq Adapters (NOVA-514151) or NEXTflex Dual-Barcode ChIP-seq Adapters with UMIs (NOVA-734103) were added at a final concentration of 0.25 µM. Libraries were then de-crosslinked at 55°C for 1hr, then 65°C overnight and cleaned with Speedbeads, 8.6% PEG and 0.8 M NaCl and eluted in TT buffer. Next, libraries were PCR amplified with 1 µM Solexa 1GA and 1GB primers and NEB Ultra II Q5 Mastermix (PCR protocol: 98°C for 30’’, then 12 or 13 cycles of 98°C for 10’’, 60°C for 15’’, 72°C for 30’’, then hold at 72°C for 2’). A final cleanup was done with Speedbeads, 8.5% PEG and 1 M NaCl, washed twice with 80% EtOH, then the final libraries were eluted in 20 µL TT. Library size was estimated by running 6 µL of eluted library on a 1.8% agarose gel and concentration was quantified with Qubit HS dsDNA kit (Invitrogen Q32854). Libraries were pooled and sequenced on Illumina NextSeq 550 or NextSeq 2000.

### Buffer components

- NP-S Buffer: 10 mM Tris-HCl pH 7.4, 0.5 mM EDTA, 50 mM NaCl, 5 mM MgCl_2_, 1 mM CaCl_2_, 0.075% NP-40, 1mM β-mercaptoethanol
- LB3: 10 mM Tris-HCl pH 7.5, 100 mM NaCl, 1 mM EDTA, 0.1% deoxycholate, 0.5% Sarkosyl
- WBI: 20 mM Tris-HCl pH 7.5, 150 mM NaCl, 1% Triton X-100, 2 mM EDTA, 1% SDS
- WBIII: 10 mM Tris-HCL pH 7.5, 250 mM LiCl, 1% Triton X-100, 1 mM EDTA, 0.7% deoxycholate
- TET: 10 mM Tris-HCl pH 8.0, 1 mM EDTA, 0.2% Tween 20
- TT Buffer: 10 mM Tris-HCl pH 8.0, 0.05% Tween 20

### Data Processing and Initial Analysis

Full details of ChIP-wrangler’s processing steps are available in the GitHub [readme](https://github.com/lapatel22/ChIP-wrangler-repo), for clarity the general steps are described below. Samples were demultiplexed with Illumina’s bcl2fastq program, adapters were removed with Trimmomatic, trimming reads once the average Phred score over 4 base pairs falls below 20. FASTQC was run for sequencing quality control. Reads were aligned with BWA MEM to a concatenated genome containing hg38 + dm6 + sacCer3. PCR duplicate reads were removed with samtools markdup, or if UMIs are present, UMIs were deduplicated with umi_tools. The alignment file was filtered to remove low quality reads (MAPQ > 50), sorted, and split by chromosome using bamUtil splitChromosome. Spike-in chromosomes were pre-labeled with a suffix (_sac3 or _dm6) before the concatenated genome was created, samtools merge was used to create species-specific alignment files, then species-specific chromosome suffixes were removed. HOMER tag directories were made; read counts for each species were filtered for primary alignments and reads were extended to model approximate ChIP-seq fragment size using the option -fragLength 150.

### Normalization

ChIP-wrangler normalization is described in detail in the GitHub [readme](https://github.com/lapatel22/ChIP-wrangler-repo), including an example where the normalization factors used in the H3K9ac titration dataset (**Fig. 1**) are calculated for reference. The spike-in normalization factor for each species is calculated as 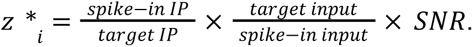. Then the normalized signal becomes: normsignal = readnorm / z*_i_. Each component of the normalization equation is described below:

#### IP enrichment over Input

IP enrichment over input is calculated as the ratio of target reads over spike-in in the IP, relative to the input. The input gives an indication of the starting levels of chromatin for each species, and the reads in the IP show how much enrichment the target epitope experiences relative to the spike-in species.

#### Estimation of IP Efficiency

Experimental factors such as inefficient or variable IP washing can lead to differences in background levels between samples, which in turn alter the proportion of the library attributable to the target epitope. This variability is often quantified as a signal-to-noise ratio using metrics like the FRiP score (Fraction of Reads in Peaks). Importantly, antibody competition between the target and the two exogenous species affects capture efficiency in both directions. For example, when the abundance of the target epitope increases, the spike-ins may show a corresponding reduction in signal-to-noise ratio. This relationship allows direct inference: if spike-in IP efficiency decreases two-fold between two samples, it implies that the target epitope levels in the second sample should be adjusted upward two-fold to reflect their relative abundance. IP efficiency is quantified at regions relevant to the given epitope of interest; e.g., IP efficiency for active histone marks such as H3K27ac and H3K4me3 is estimated by quantifying around the peak of signal, from -100 to +700bp of the TSS for each species. Additional details are provided in the “Estimate IP Efficiency” section of the Github readme.

#### Normalization Factor Scaling

Normalization factors for each IP can be scaled in two ways, so that spike-in factors remain in comparable dynamic ranges:

1. **Average scaling within each treatment group,** or
2. **Normalization relative to a reference control**

Method (1) is agnostic to experimental conditions and avoids the need to explicitly select a reference. However, normalization factors then depend on all samples in the experiment: a single sample with an unusually high factor (>> 1) will drive other samples artificially low (<< 1), compressing their data. In contrast, method (2) where samples are normalized to a reference control is more likely to preserve biological relevance. Additional details and example data are provided in the “Get Sequencing Stats” section of the Github readme.

From above, each normalization factor for a given sample is calculated by:

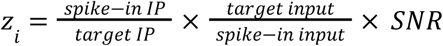

Letting Z = {z_1_, z_2_, z_3_, …, z_n_} be the set of normalization factors in an experiment, where *C* ⊆ *Z* is the set of normalization factors corresponding to control samples:

Method 1 - Experiment-wide Mean Scaling

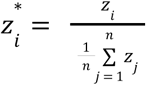

Method 2 - Control-only Mean Scaling

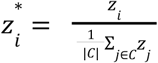

#### Dual Spike-ins

Each sample is normalized to spike-ins *D. melanogaster* and *S. cerevisiae* separately, then the average of each normalization factor is used for dual normalization. For each sample, bedGraph files and HOMER Tag Directories of signals normalized by each method are created.

The accuracy of each normalization method is quantified by calculating the R^2^ value of data against the line of expected signal (diagonal line shown in plots of **Fig. 1c**). The line of expected signal is simply the diagonal line connecting maximum and minimum signal. R^2^ is calculated by the equation 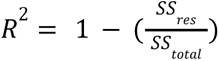 where 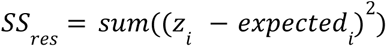 and 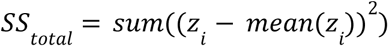. R^2^ values are reported in **Fig. 1c** and **Fig. 3c** as inserts in plots.

#### Spike-in Normalization Factor Delta

To estimate the differential between normalization factors generated from each spike-in species, the spike-in normalization factor delta is calculated as the difference between each spike-in normalization factor from the average normalization factor, divided by the average normalization factor. As the normalization factor is a dividing factor (normsignal = readnorm / z*_i_), the *negative delta* in **Fig. 2c** is plotted for clarity (here, a value above zero results in an increase in normalized signal in **Fig. 2d**). Normalization factors derived from each spike-in are expected to correlate, ChIP-wrangler’s workflow includes a QC report of each species’ normalization factor, the dual normalization factor, and flags if the two species’ normalization factors are too dissimilar. More details are in the section “QC Report” in the readme.

### Visualization

Promoter histograms were generated with HOMER annotatePeaks with the option -size 4000 -hist 25 at RefSeq TSSs in **Figs 1, 3 and 4**. Gene body histograms are generated with HOMER annotatePeaks -from -500 to 50000 -hist 100 as in **Supp Fig. 7**. Histogram confidence intervals created by plotting the variation in the results of single spike-in species normalization. Dot plots of normalized signal in HeLa-S3 H3K9ac (**Fig. 1**) and H3K27ac (**Fig. 3**) titration datasets were generated by quantifying the signal at promoters or peaks using area under the curve (see Github tutorial for examples, source metagene data in **Fig. 1c**, **Supp. Fig. 4**). Nucleotide frequency plots were created using tagFreqUniq.txt files generated by HOMER makeTagDirectory using the option -checkGC and -fragLength 150. Scatterplots were generated from HOMER annotatePeaks quantification of signal at peak / TSS regions or genomic bins as specified in figure captions. Peak finding for histone marks done with HOMER findPeaks using options -style histone -size 1000 -minDist 2500. When fold changes are plotted (violin or scatterplot), data are first filtered to remove low-intensity signals below the mean signal in the untreated condition. When specified, quantiling of RPB1 signal is done by first averaging signals in untreated control samples, then splitting data into 4 quantiles based on average untreated signal intensity. To compare Triptolide and auxin treatments, HOMER findPeaks was run on each sample, then peaks in each condition were merged to find a common set of peaks for each histone mark. Levels of the mark are then quantified at common regions.

For differential ChIP-seq peak analysis, a raw counts matrix of H3K27ac signal was generated with HOMER annotatePeaks on the merged peak set with option -raw. Default size factors used by DESeq2 were estimated with getSizeFactors on the dds object, which were multiplied by the dual spike-in normalization factor. Then DESeq2 was run with these custom normalization factors to identify differential peaks. Thresholds of apeglm-shrunken Log2FC > 1, FDR < 0.05 for upregulated peaks and apeglm-shrunken Log2FC < -1 FDR < 0.05 for downregulated peaks were used.

Putative enhancers were determined by finding distal H3K27ac peaks (further than 3kb from RefSeq TSSs), then H3K4me1 and RPB1 signal was quantified at distal peaks (**Supp. Fig. 7**). Whenever specified, promoter-distal regions refer to all regions further than 3kb from RefSeq TSSs.

Plots generated with R, ggPlot2, matplotlib, and Illumina Sequencing Analysis Viewer 3.0.0. Additional details and code used to generate figures available on [Github](https://github.com/lapatel22/ChIP-wrangler-repo/).

## Data Availability

Previously published data available at GSE273915. Data generated for this manuscript is available at GSE286875. Details and tutorial for ChIP-wrangler and code used to generate figures are available on [Github](https://github.com/lapatel22/natgen_presubmit_2024/tree/master). UCSC Genome Browser sessions are provided as links within the caption of the relevant figures, static browser images are included as Supplemental Figures. Raw immunoblot images are available in Supplemental Material.

