## Supplementary material for "Improved spike-in normalization clarifies the relationship between active histone modifications and transcription": patel-etal-biorxiv-sup-figs-revision2-submit

**Table of Contents**

**Supp. Fig. 1. Additional Data and QCs for Fig. 1, a titration of H3K9ac levels in HeLa-S3 cells.**

**Supp. Fig. 2. Normalizing to dual spike-ins modest increases in accuracy compared to each species individually and dramatically compared to read-depth normalization.**

**Supp. Fig. 3. UCSC genome browser tracks for Fig. 1, a titration of H3K9ac levels in HeLa-S3 cells.**

**Supp. Fig. 4. Additional data and QCs for Fig. 2, an example of the increased rigor obtained by using spike-ins from two species.**

**Supp. Fig. 5. UCSC genome browser tracks for Fig. 2, an example of the increased rigor obtained by using spike-ins from two species.**

**Supp. Fig. 6. Additional data for Fig. 3, a titration of H3K27ac levels and spike-in chromatin in HeLa-S3 cells.**

**Supp. Fig. 7. UCSC genome browser tracks for Fig. 3, a titration of H3K27ac levels and spike-in chromatin in HeLa-S3 cells.**

**Supp. Fig. 8. ChIP-seq using Rpb1 antibody does not show enrichment for RnapII in S. cerevisiae.**

**Supp. Fig. 9. ChIP-wrangler normalized H3K4me3 at M. musculus RefSeq promoters after degradation of COMPASS subunits.**

**Supp. Fig. 10. Additional Data for Fig. 4, Changes in H3K36me3, H3K4me1 and Rpb1 after Triptolide and auxin.**

**Supp. Fig. 11. Additional data for Fig. 4, changes in H3K27ac, H3K4me3, and Rpb1 after treatment of HCT116-RPB1 degron cells with Triptolide and auxin.**

**Supp. Fig. 12. UCSC genome browser tracks for Fig. 4, treatment of HCT116-RPB1 degron cells with Triptolide and auxin.**

**Supp. Fig. 13. Additional data for Fig. 4, spike-in data and QC metrics for treatment of HCT116-RPB1 degron cells with Triptolide and auxin.**

**Supp. Fig. 14. Additional data for Figure 5, spike-in QC metrics for HeLa-S3 Rpb1 data.**

**Supp Fig 15. Pairwise comparisons of changes in H3K27ac at genomic regions using Wilcoxon rank sum test.**

**Supp. Fig. 16. Differential H3K27ac Peak Finding after Triptolide or auxin-mediated degradation of Rpb1.**

**Supp Fig. 17. UCSC genome browser tracks for additional biological replicates of treatment of HCT116-RPB1 degron cells with Triptolide an auxin.**

**Supp. Fig. 18. Examples of expected quality metric results provided by ChIP-wrangler.**

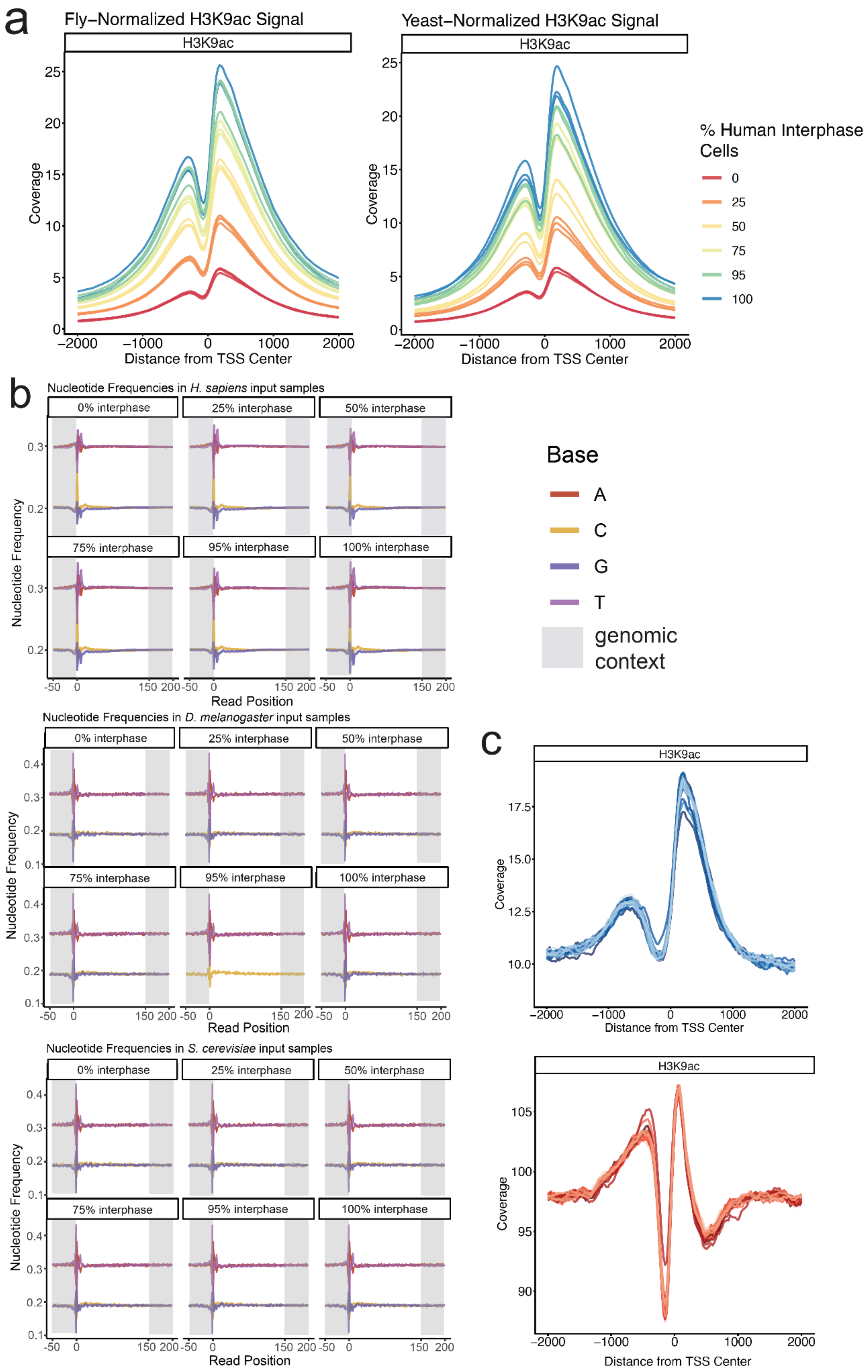

a
Fly–Normalized H3K9ac Signal
Yeast–Normalized H3K9ac Signal
H3K9ac
Coverage
Distance from TSS Center
% Human Interphase Cells
0
25
50
75
95
100
b
Nucleotide Frequencies in H. sapiens input samples
Nucleotide Frequencies in D. melanogaster input samples
Nucleotide Frequencies in S. cerevisiae input samples
0% interphase
25% interphase
50% interphase
75% interphase
95% interphase
100% interphase
Nucleotide Frequency
Read Position
Base
A
C
G
T
genomic context
c
H3K9ac
Coverage
Distance from TSS Center

**Supp. Fig. 1. Additional Data and QCs for Fig. 1, a titration of H3K9ac levels in HeLa-S3 cells.**

**a.** Histogram of *H. sapiens* H3K9ac signal at RefSeq TSSs normalized to either *D. melanogaster* (left) or *S. cerevisiae* (right).

**b.** Nucleotide frequency plots of reads aligned to *H. sapiens*, *D. melanogaster* and *S. cerevisiae* in input libraries.

**c.** Histogram of H3K9ac at *D. melanogaster* or *S. cerevisiae* RefSeq TSSs.

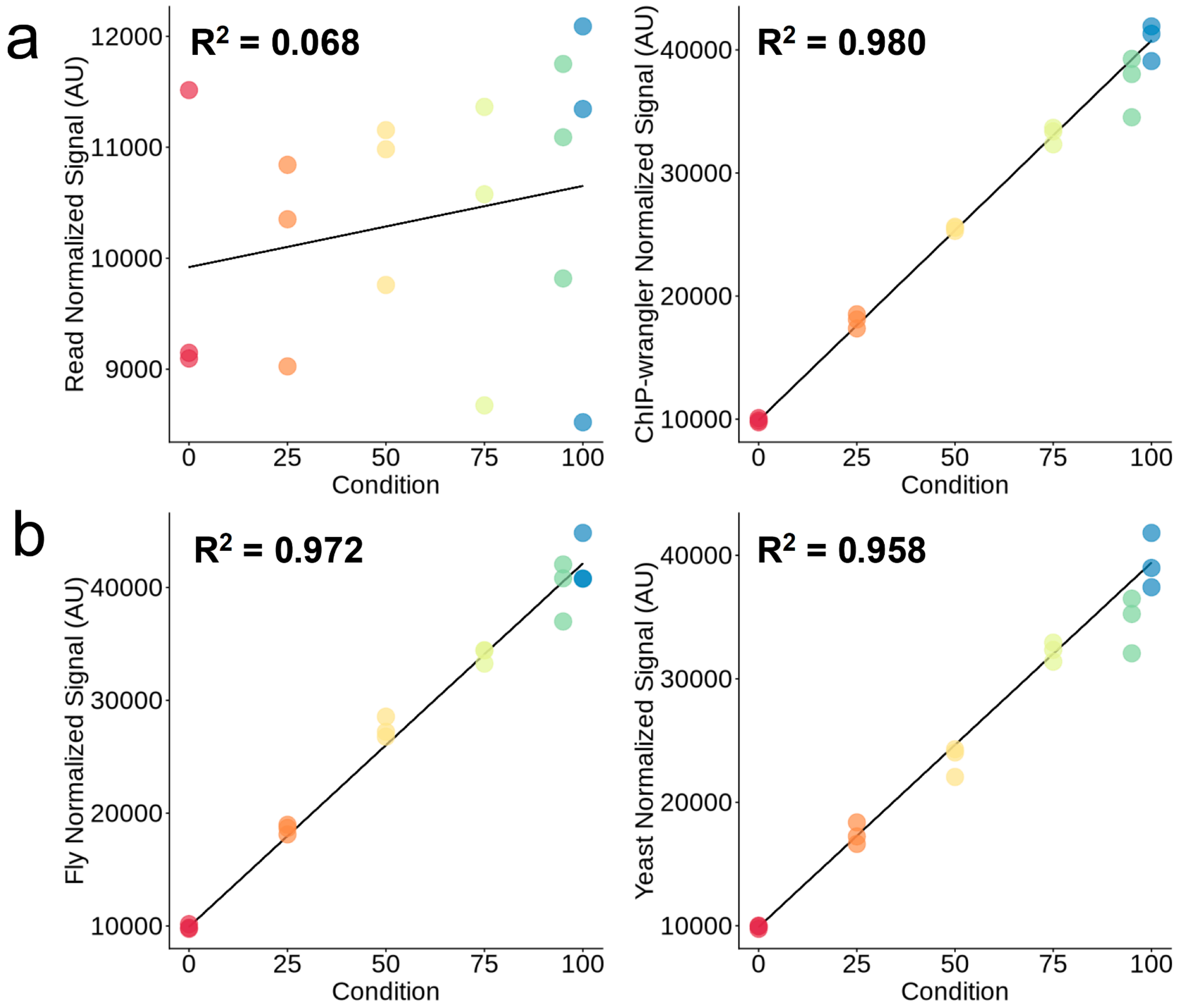


**Supp. Fig. 2. Normalizing to dual spike-ins modest increases in accuracy compared to each species individually and dramatically compared to read-depth normalization.**

**a.** Adapted from **Fig. 1d**, total H3K9ac signal is quantified at promoters after read-depth normalizing (left) or dual spike-in normalization with ChIP-wrangler.

**b.** Total H3K9ac signal is quantified at promoters after normalizing to either *D. melanogaster* (left) or *S. cereivisae* (right). Accuracy of each normalization method is calculated by $R^2$, in plot inserts.

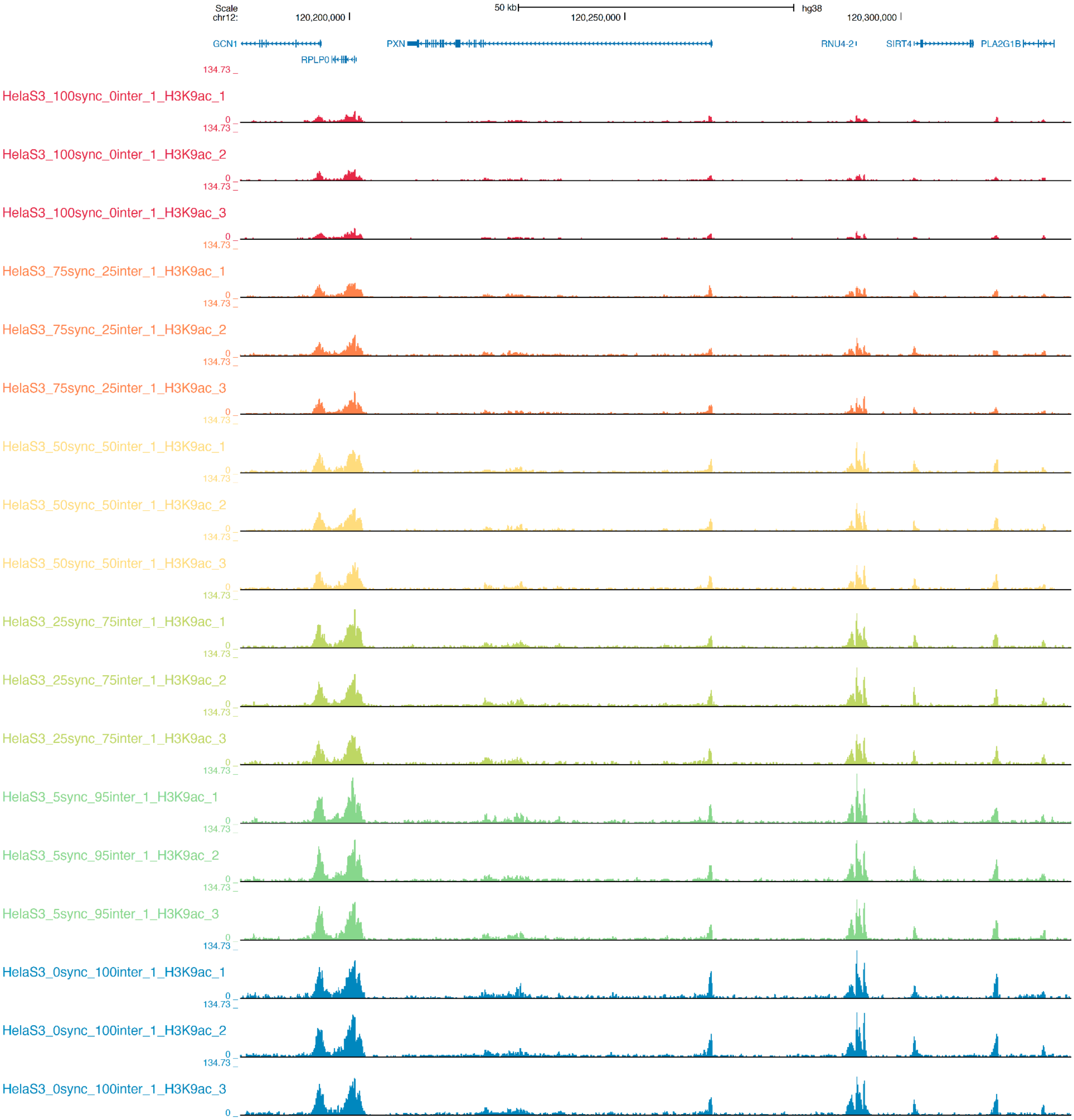


**Supp. Fig. 3. UCSC genome browser tracks for Fig. 1, a titration of H3K9ac levels in HeLa-S3 cells.** ChIP-wrangler normalized H3K9ac from a titration of low to high signal generated by mixing mitotic-arrested and unsynchronized (interphase) HeLa-S3 cells.

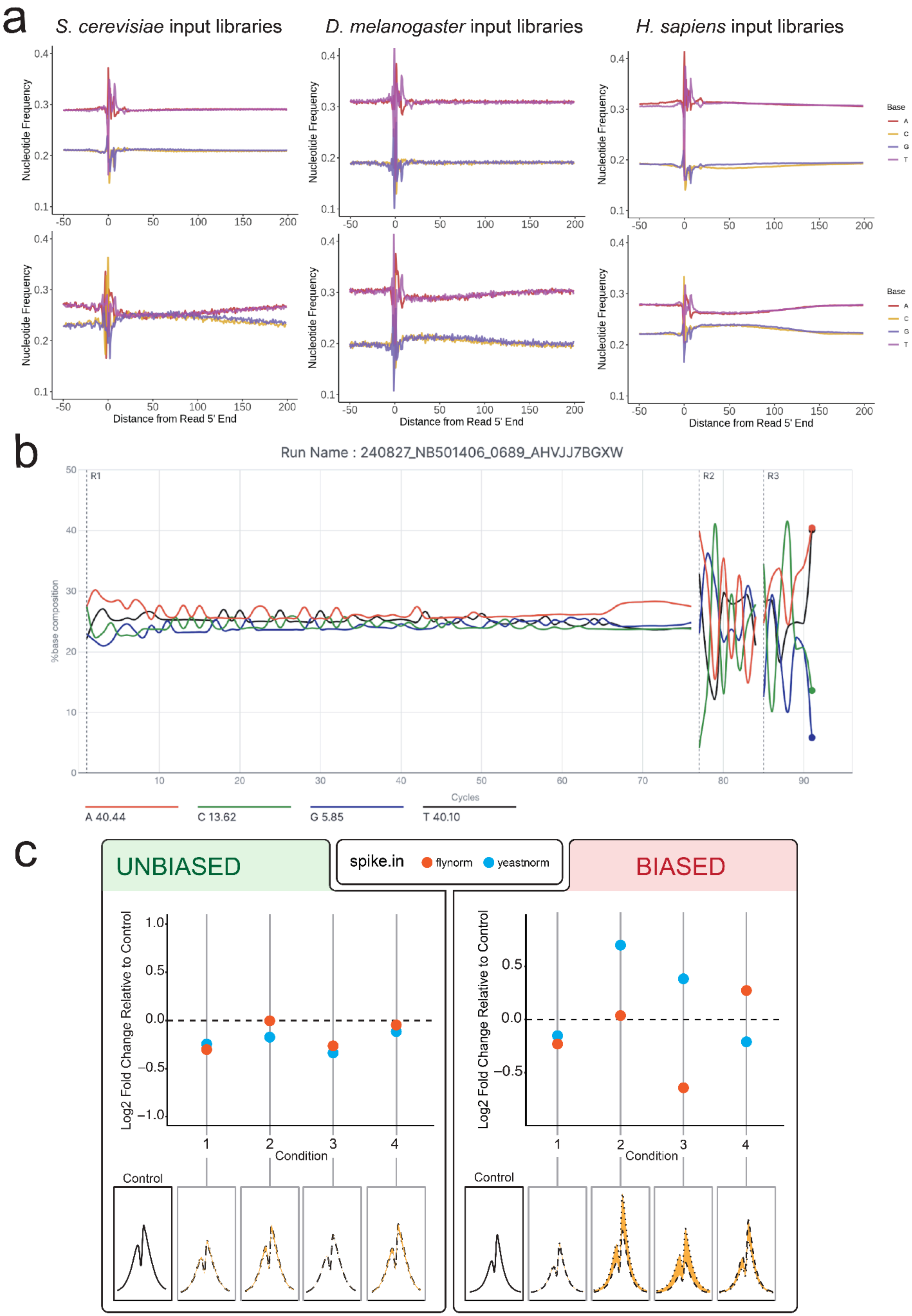

a
S. cerevisiae input libraries
D. melanogaster input libraries
H. sapiens input libraries
Nucleotide Frequency
Distance from Read 5' End
Base
A
C
G
T
b
Run Name : 240827_NB501406_0689_AHVJJ7BGXW
%base composition
Cycles
R1
R2
R3
A 40.44
C 13.62
G 5.85
T 40.10
c
UNBIASED
spike.in
flynorm
yeastnorm
BIASED
Log2 Fold Change Relative to Control
Condition
Control

**Supp. Fig. 4. Additional data and QCs for Fig. 2, an example of the increased rigor obtained by using spike-ins from two species.**

**a.** Nucleotide frequency plots for the unbiased (top) and biased ChIP-seq input libraries (bottom) for *H. sapiens*, *D. melanogaster*, and *S. cerevisiae* (left to right).

**b.** The nucleotide frequency distribution of the Illumina Nextseq500 run used in the biased experiment. Generated with Illumina Sequencing Analysis Viewer 3.0.0. Given that this plot is generated for a pool of input and IP-enriched libraries, the GC bias was not immediately observable.

**c.** Log2 Fold Change of H3K27ac in each treatment condition relative to control, in unbiased (left) and biased (right) experiments. Metagene plots of H3K27ac signal at TSS used to calculate overall change in signal in the dot plots are shown below the graph. The shaded orange area represents the difference in result when normalizing to each spike-in species.

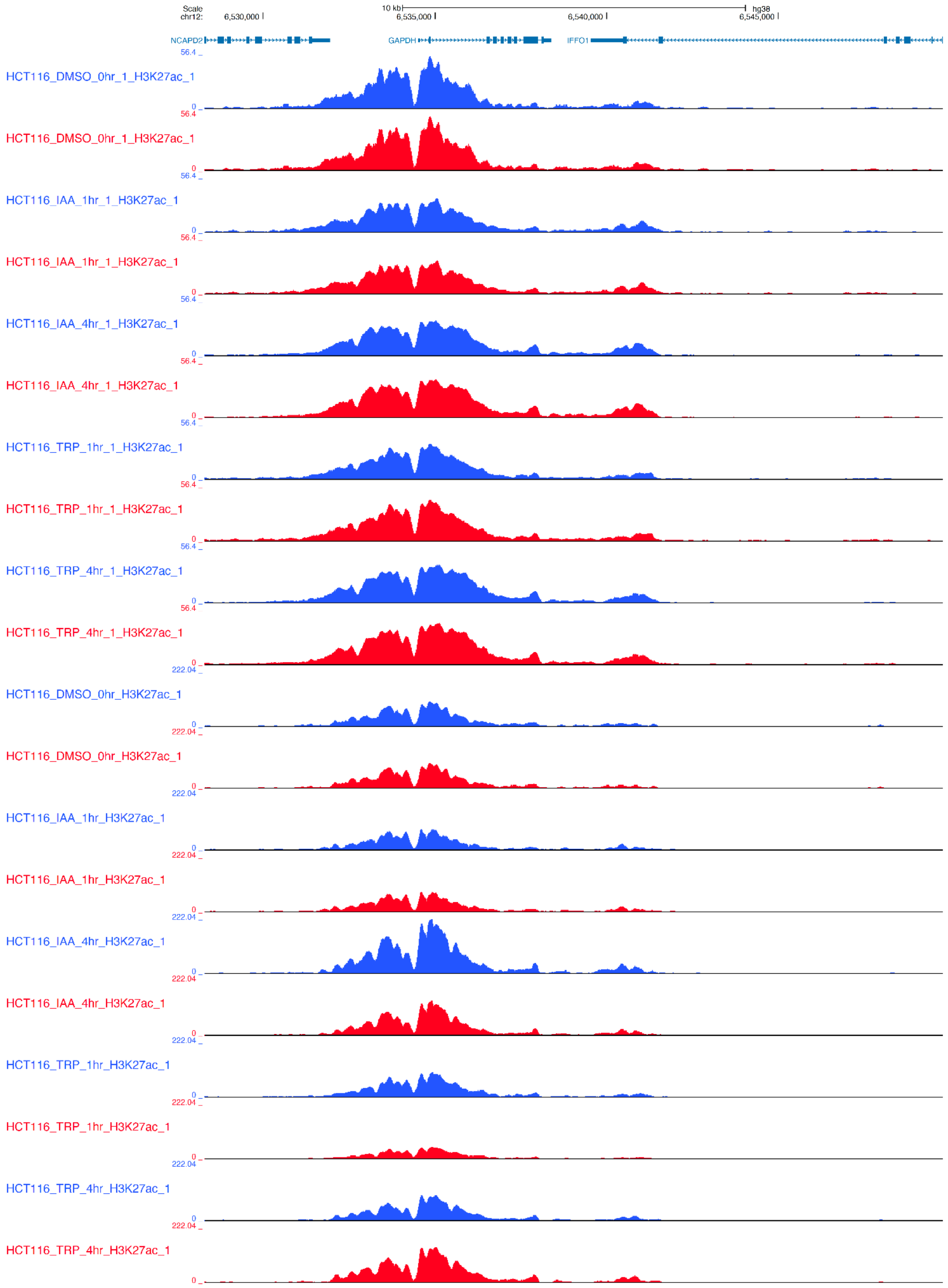
Scale
chr12:
10 kb
hg38
6,530,000
6,535,000
6,540,000
6,545,000
NCAPD2
GAPDH
IFFO1
HCT116_DMSO_0hr_1_H3K27ac_1
HCT116_DMSO_0hr_1_H3K27ac_1
HCT116_IAA_1hr_1_H3K27ac_1
HCT116_IAA_1hr_1_H3K27ac_1
HCT116_IAA_4hr_1_H3K27ac_1
HCT116_IAA_4hr_1_H3K27ac_1
HCT116_TRP_1hr_1_H3K27ac_1
HCT116_TRP_1hr_1_H3K27ac_1
HCT116_TRP_4hr_1_H3K27ac_1
HCT116_TRP_4hr_1_H3K27ac_1
HCT116_DMSO_0hr_H3K27ac_1
HCT116_DMSO_0hr_H3K27ac_1
HCT116_IAA_1hr_H3K27ac_1
HCT116_IAA_1hr_H3K27ac_1
HCT116_IAA_4hr_H3K27ac_1
HCT116_IAA_4hr_H3K27ac_1
HCT116_TRP_1hr_H3K27ac_1
HCT116_TRP_1hr_H3K27ac_1
HCT116_TRP_4hr_H3K27ac_1
HCT116_TRP_4hr_H3K27ac_1
56.4
222.04
0

**Supp. Fig. 5. UCSC genome browser tracks for Fig. 2, an example of the increased rigor obtained by using spike-ins from two species.** ChIP-wrangler normalized H3K27ac in HCT116-RPB1 degron cells treated with Triptolide or auxin for 1 and 4 hours. Each sample is normalized separately to *D. melanogaster* (blue) or *S. cerevisiae* (red). Top: unbiased libraries show consistent normalization to both spike-in species; bottom: normalization to *D. melanogaster* or *S. cerevisiae* show result in different levels of H3K27ac.

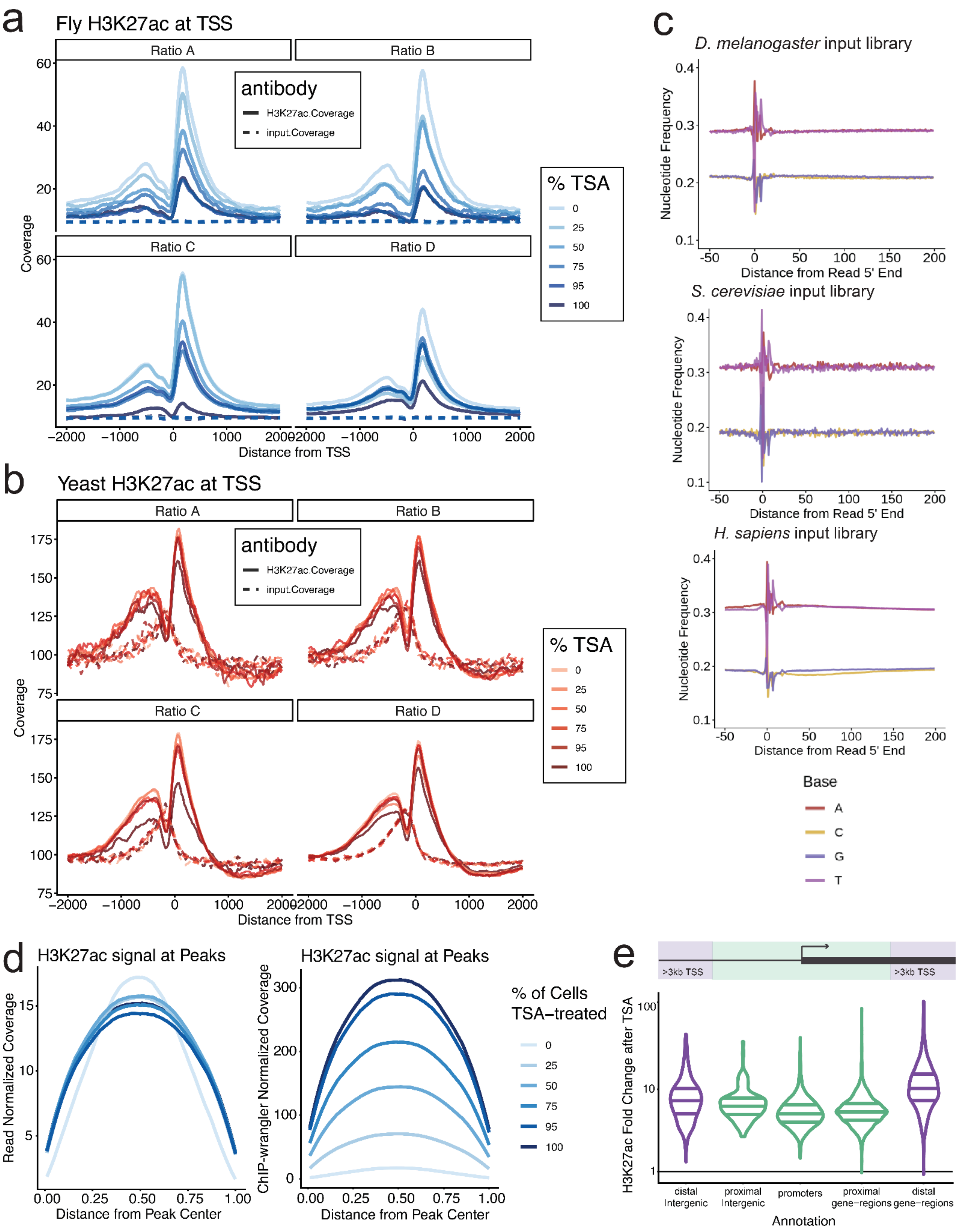
a
Fly H3K27ac at TSS
Ratio A
Ratio B
Ratio C
Ratio D
antibody
H3K27ac.Coverage
input.Coverage
% TSA
0
25
50
75
95
100
Coverage
Distance from TSS
b
Yeast H3K27ac at TSS
c
D. melanogaster input library
S. cerevisiae input library
H. sapiens input library
Nucleotide Frequency
Distance from Read 5' End
Base
A
C
G
T
d
H3K27ac signal at Peaks
Read Normalized Coverage
ChIP-wrangler Normalized Coverage
Distance from Peak Center
% of Cells TSA–treated
e
>3kb TSS
H3K27ac Fold Change after TSA
distal Intergenic
proximal Intergenic
promoters
proximal gene−regions
distal gene−regions
Annotation

f

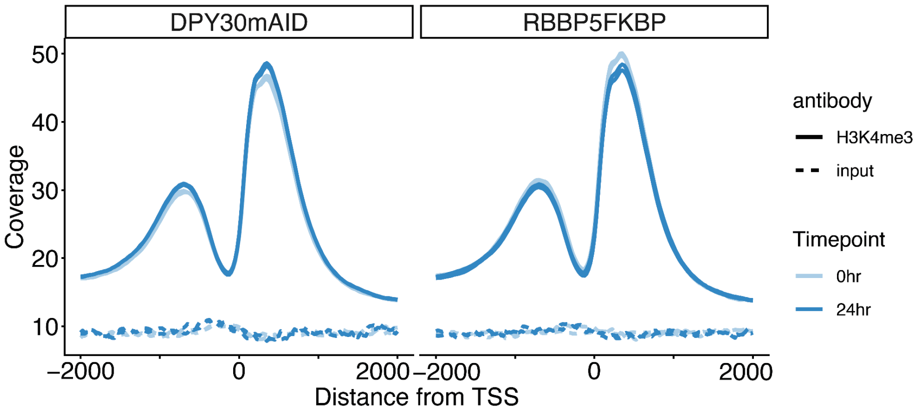


g

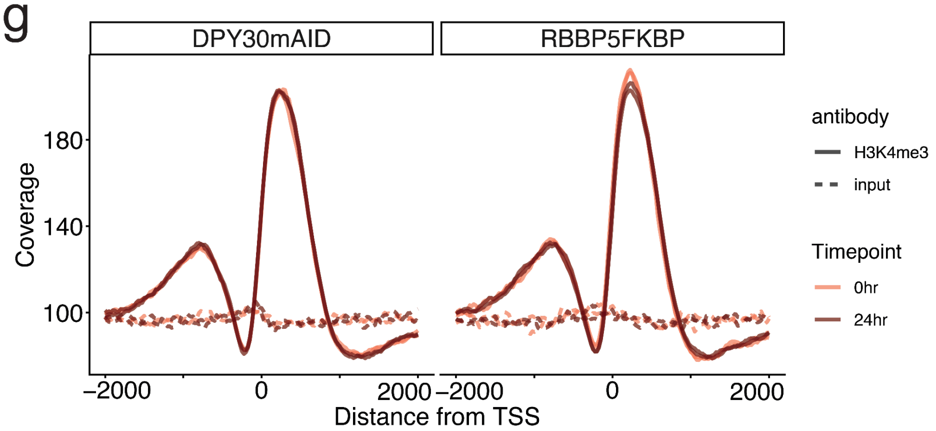


**Supp. Fig. 6. Additional data for Fig. 3, a titration of H3K27ac levels and spike-in chromatin in HeLa-S3 cells.**

**a.** Histogram of H3K27ac and input signal at *D. melanogaster* RefSeq TSSs.

**b.** Histogram of H3K27ac and input signal at *S. cerevisiae* RefSeq TSSs.

**c.** Representative nucleotide frequency plots in ChIP-seq inputs.

**d.** Metagene plot of *H. sapiens* H3K27ac signal at merged peaks from Ratio B, either read-depth normalized (left) or ChIP-wrangler normalized (right).

**e.** Fold change in ChIP-wrangler normalized H3K27ac signal after 1μM TSA-treatment at all peaks, binned by genomic annotation and promoter proximity.

**f.** Metagene plot of H3K4me3 at *D. melanogaster* RefSeq TSSs in mESC-Dpy30mAID and mESC-Rbbp5FKBP degradation datasets.

**g.** Metagene plot of H3K4me3 at *S. cerevisiae* RefSeq TSSs in mESC-Dpy30mAID and mESC-Rbbp5FKBP degradation datasets.

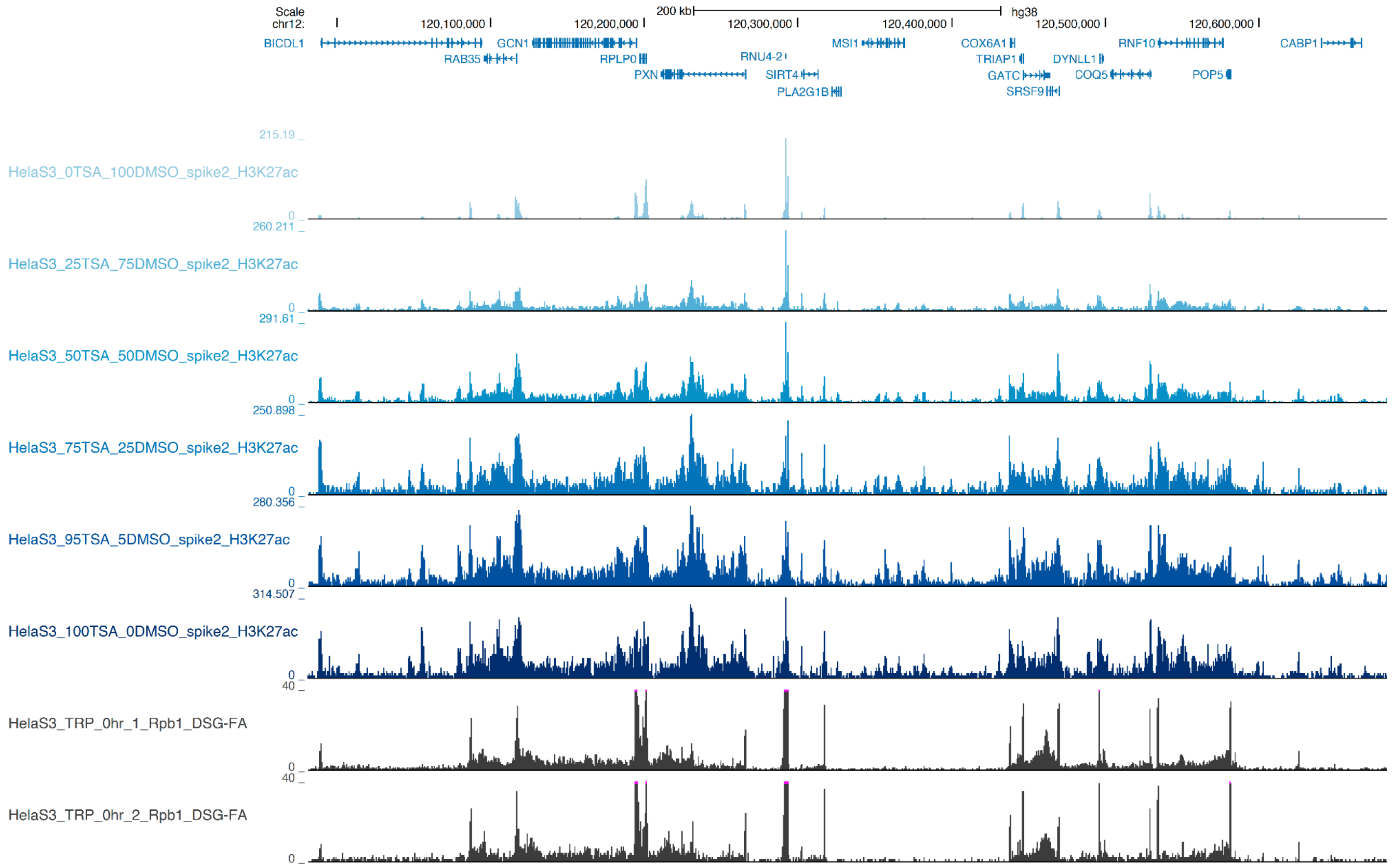


**Supp. Fig. 7. UCSC genome browser tracks for Fig. 3, a titration of H3K27ac levels and spike-in chromatin in HeLa-S3 cells.**

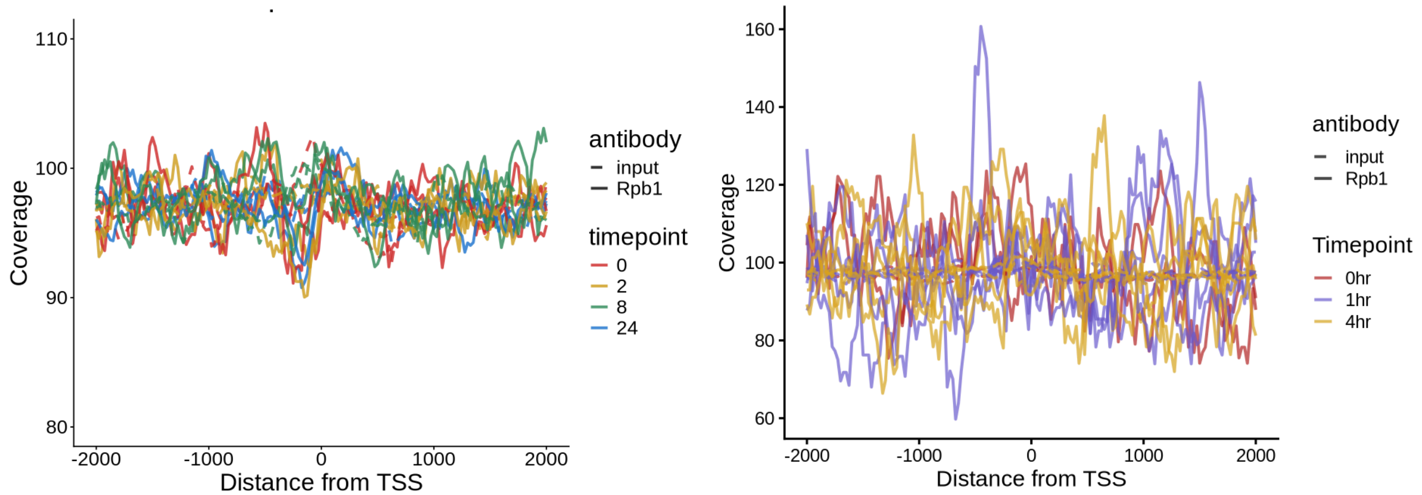


**Supp. Fig 8. ChIP-seq using Rpb1 antibody does not show enrichment for RnapII in *S. cerevisiae*.**
**Left:** Histogram of Rpb1 (Rpb1-NTD, Cell Signaling Technologies Cat #14958) at RefSeq promoters, *S. cerevisiae* was added as a spike-in to mESC-Dpy30-mAID and mESC-Rbbp5-FKBP lysates for ChIP-seq. **Right:** Histogram of Rpb1 at *S. cerevisiae* around RefSeq promoters, *S. cerevisiae* was added as a spike-in to HCT116-RPB1 degron lysates. Dashed lines indicate input samples, solid lines are Rpb1 ChIP-seq samples.

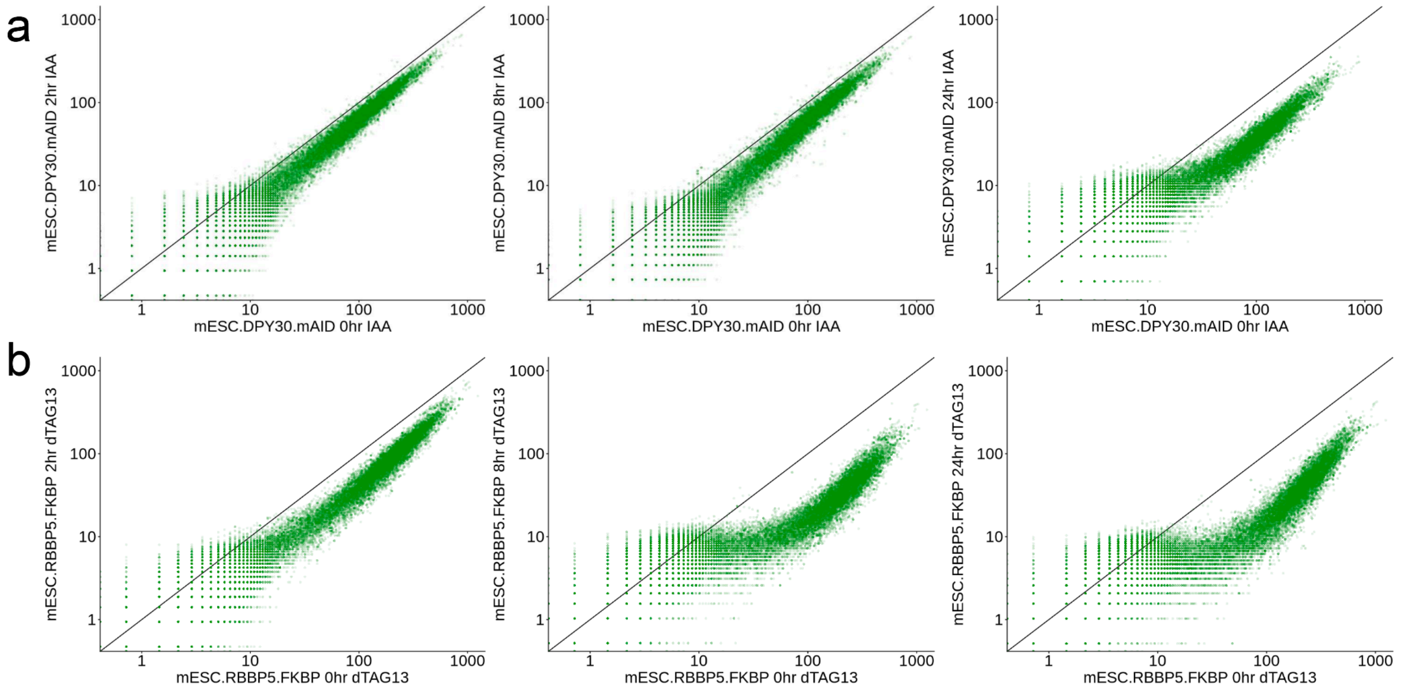


**Supp. Fig. 9. ChIP-wrangler normalized H3K4me3 at *M. musculus* RefSeq promoters after degradation of COMPASS subunits.**

**a.** Scatterplot of H3K4me3 signal at promoters in mESC-Dpy30-mAID samples after treatment with IAA for 2, 8, or 24 hours compared to control (left to right).

**b.** Scatterplot of H3K4me3 signal at promoters in mESC-Rbbp5-FKBP samples after treatment with dTAG-13 for 2, 8, or 24 hours compared to control (left to right).

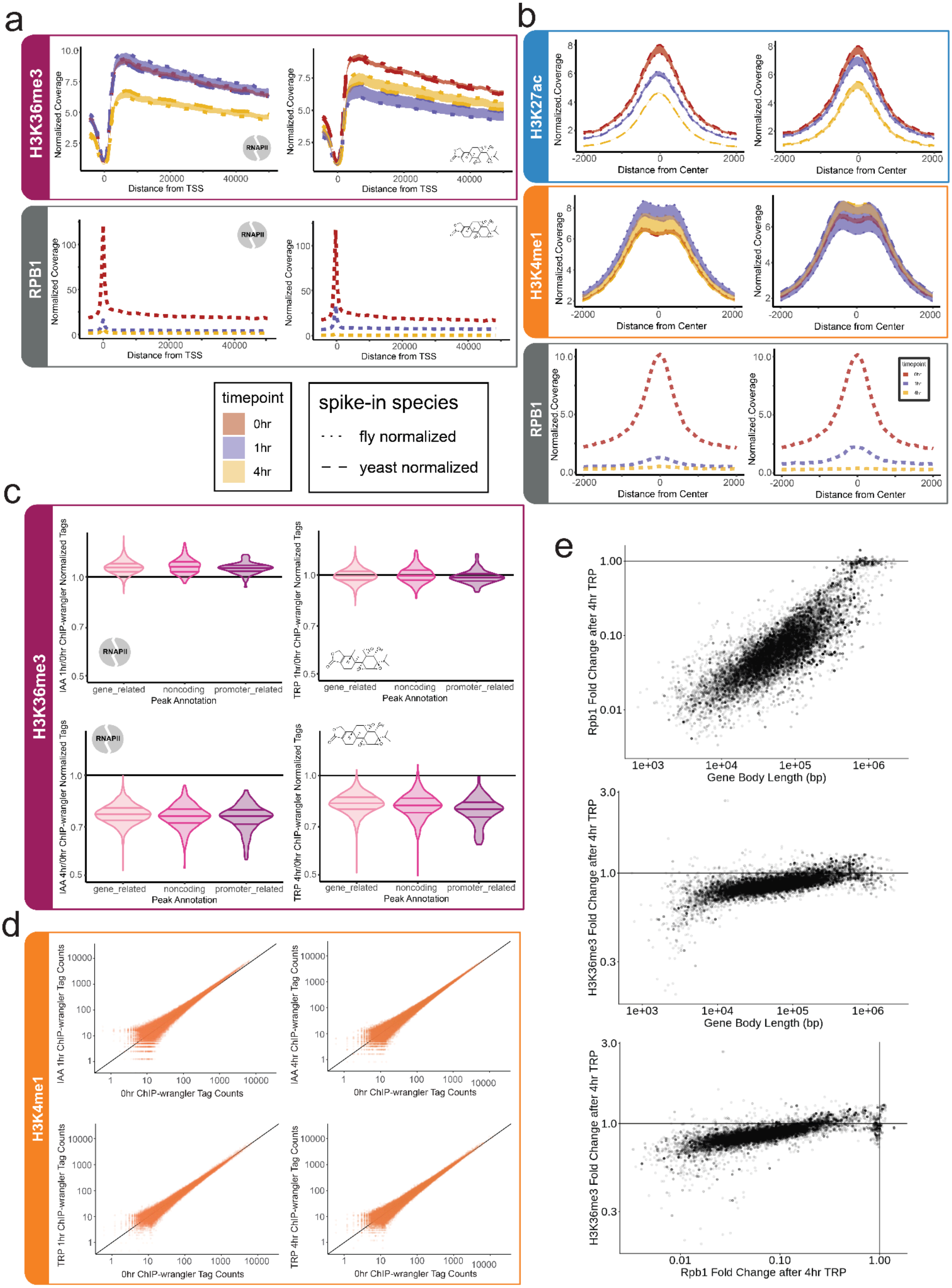

a
H3K36me3
RPB1
Normalized.Coverage
Distance from TSS
RNAPII
Normalized Coverage
timepoint
0hr
1hr
4hr
spike-in species
fly normalized
yeast normalized
b
H3K27ac
H3K4me1
RPB1
Normalized.Coverage
Distance from Center
timepoint
c
H3K36me3
IAA 1hr/0hr ChIP-wrangler Normalized Tags
TRP 1hr/0hr ChIP-wrangler Normalized Tags
IAA 4hr/0hr ChIP-wrangler Normalized Tags
TRP 4hr/0hr ChIP-wrangler Normalized Tags
gene_related
noncoding
promoter_related
Peak Annotation
d
H3K4me1
IAA 1hr ChIP-wrangler Tag Counts
IAA 4hr ChIP-wrangler Tag Counts
TRP 1hr ChIP-wrangler Tag Counts
TRP 4hr ChIP-wrangler Tag Counts
0hr ChIP-wrangler Tag Counts
e
Rpb1 Fold Change after 4hr TRP
Gene Body Length (bp)
H3K36me3 Fold Change after 4hr TRP

**Supp. Fig. 10. Additional Data for Fig. 4, Changes in H3K36me3, H3K4me1 and RPB1 after Triptolide and auxin.**

**a.** Histograms of ChIP-wrangler normalized H3K36me3 and RPB1 signal along gene bodies after treatment of HCT116-RPB1 degron cells with Triptolide or auxin.

**b.** Histograms of ChIP-wrangler normalized H3K27ac, H3K4me1, and RPB1 signal around putative enhancers, defined as H3K27ac peaks > 3kb from a TSS.

**c.** Fold Change in H3K36me3 after treatment with auxin or Triptolide, binned by genome annotation.

**d.** Scatterplot of ChIP-wrangler normalized H3K4me1 signal of Triptolide and auxin treatments compared to untreated controls at merged H3K4me1 peaks.

**e.** Changes in RPB1 and H3K36me3 after 4 hours of Triptolide at active genes (top 50% of untreated RPB1 signal), relative to gene body length. Bottom: fold change in H3K36me3 versus fold change in RPB1 at active genes after 4 hours of Triptolide.

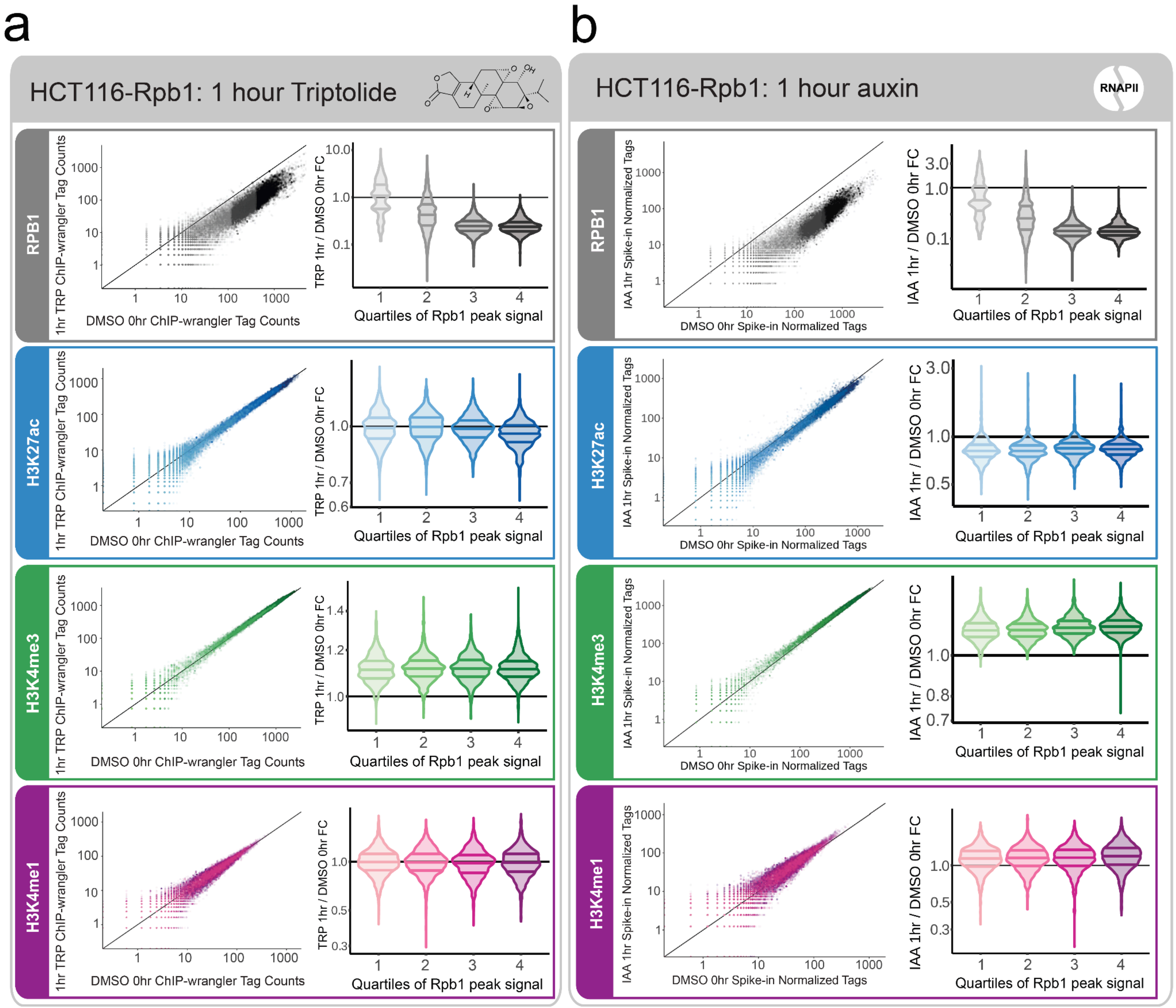


**Supp. Fig. 11. Additional data for Fig. 4, changes in H3K27ac, H3K4me3, and RPB1 after treatment of HCT116-RPB1 degron cells with Triptolide and auxin.**

**a.** Comparison of RPB1, H3K27ac, H3K4me3, and H3K4me1 levels at promoters after 1 hour of auxin relative to untreated levels in HCT116-RPB1 degron cells, shaded according to the quartiles of untreated RPB1 signal, displayed as violin plots on the right.

**b.** Comparison of RPB1, H3K27ac, H3K4me3, and H3K4me1 levels at promoters after 1 hour of auxin relative to untreated levels in HCT116-RPB1 degron cells, shaded according to the quartiles of untreated RPB1 signal, displayed as violin plots on the right.

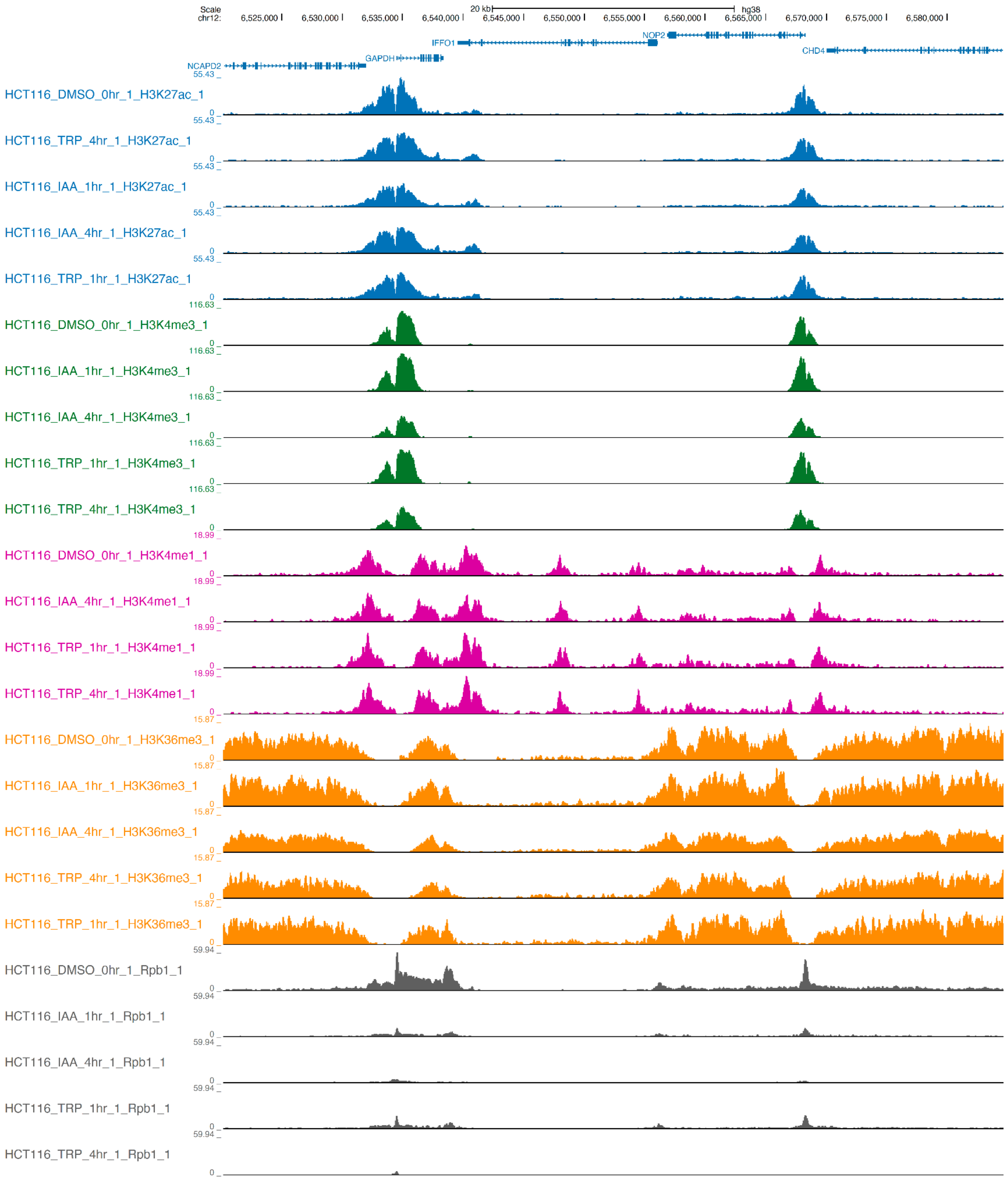


**Supp. Fig. 12. UCSC genome browser tracks for Fig. 4, treatment of HCT116-RPB1 degron cells with Triptolide and auxin.** Cells were treated with either Triptolide or auxin for 1 and 4 hours, then ChIP-seq was performed for H3K27ac, H3K4me3, H3K4me1, H3K36me3, and RPB1. All data are normalized with ChIP-wrangler, histone marks were normalized to dual spike-ins *D. melanogaster* and *S. cerevisiae*, while RPB1 data were normalized to *D. melanogaster*.

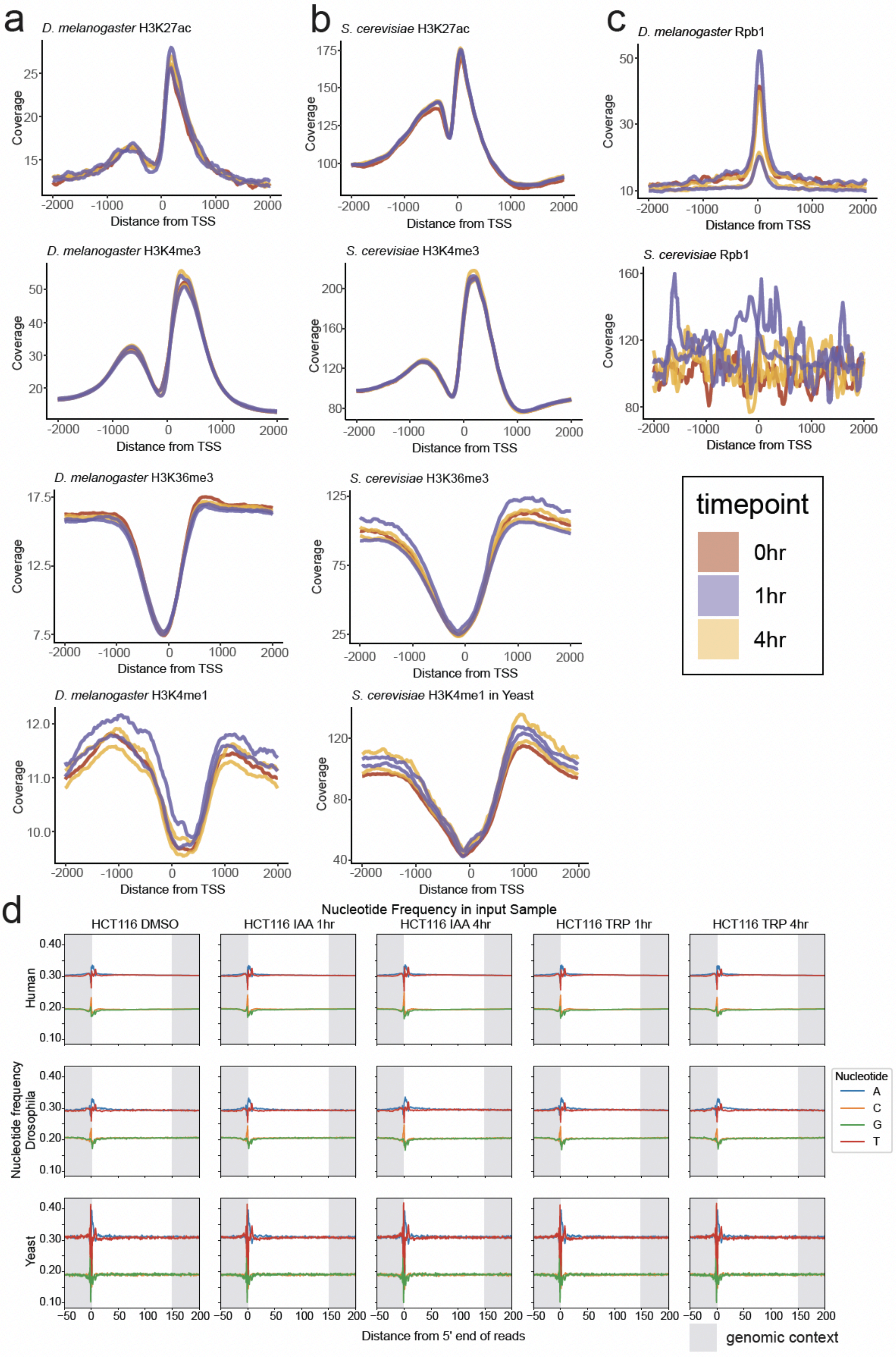
a
D. melanogaster H3K27ac
D. melanogaster H3K4me3
D. melanogaster H3K36me3
D. melanogaster H3K4me1
b
S. cerevisiae H3K27ac
S. cerevisiae H3K4me3
S. cerevisiae H3K36me3
S. cerevisiae H3K4me1 in Yeast
c
D. melanogaster Rpb1
S. cerevisiae Rpb1
Coverage
Distance from TSS
timepoint
0hr
1hr
4hr
d
Nucleotide Frequency in input Sample
HCT116 DMSO
HCT116 IAA 1hr
HCT116 IAA 4hr
HCT116 TRP 1hr
HCT116 TRP 4hr
Human
Drosophila
Yeast
Nucleotide frequency
Nucleotide
A
C
G
T
Distance from 5' end of reads
genomic context

**Supp. Fig. 13. Additional data for Fig. 4, spike-in data and QC metrics for treatment of HCT116-RPB1 degron cells with Triptolide and auxin. a.** Histograms of histone modification ChIP-seq signal at *D. melanogaster* RefSeq TSSs. **b.** Histograms of histone modification ChIP-seq signal at *S. cerevisiae* RefSeq TSSs. **c.** Histograms of Rpb1 signal in *D. melanogaster* and *S. cerevisiae* at RefSeq TSSs. **d.** Nucleotide Frequency plots of *H. sapiens*, *D. melanogaster*, and *S. cerevisiae* aligned reads for all input samples.

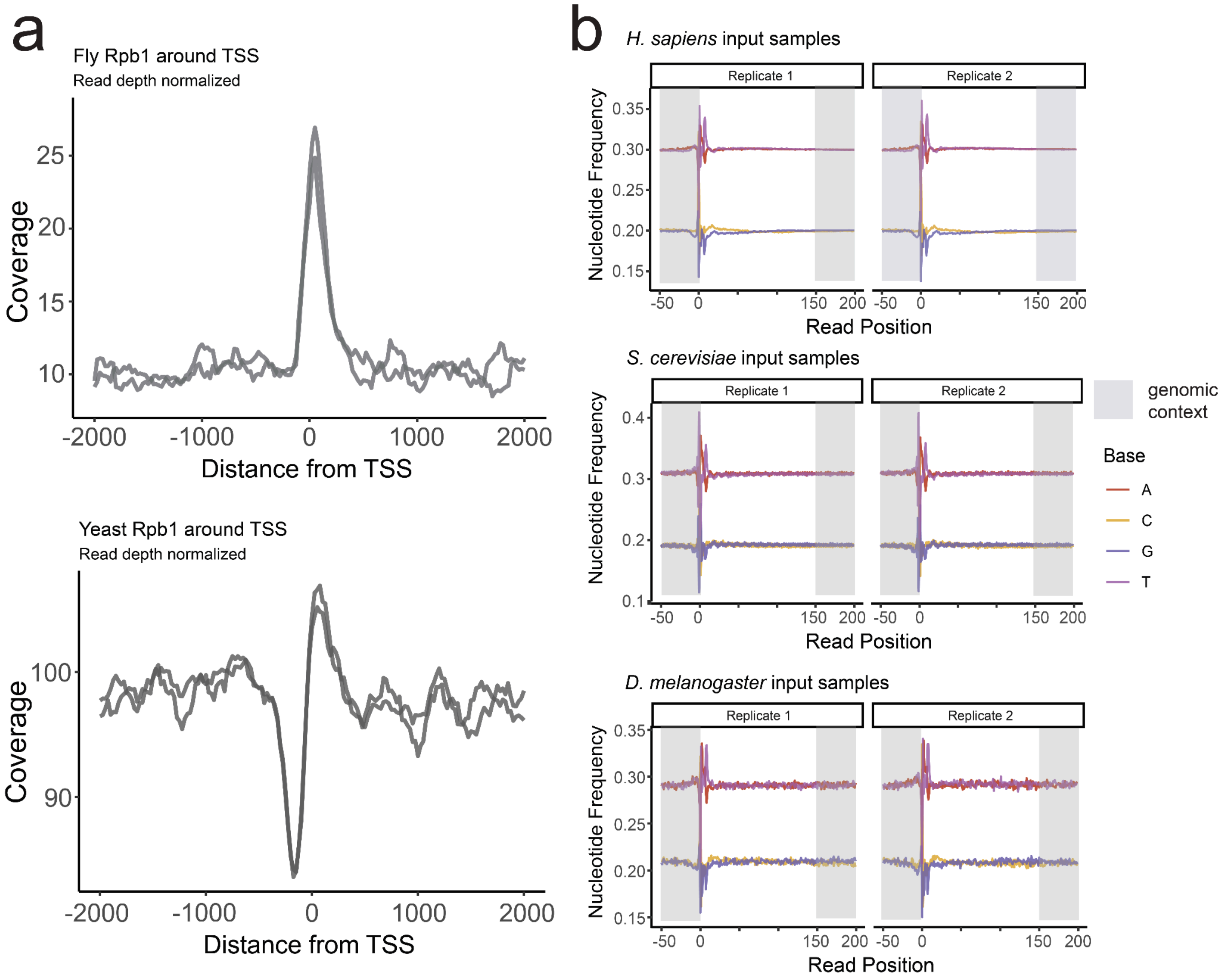


**Supp. Fig. 14. Additional data for Figure 5, spike-in QC metrics for HeLa-S3 Rpb1 data.**

**a.** Histograms of Rpb1 ChIP-seq signal at *D. melanogaster* RefSeq TSSs (top), and at *S. cerevisiae* RefSeq TSSs (bottom).

**b.** Nucleotide Frequency plots of *H. sapiens*,*S. cerevisiae,* and *D. melanogaster* aligned reads for input samples.

| | Intergenic Distal | Intergenic Proximal | Promoter | Proximal Gene Body | Distal Gene Body |
|---|---|---|---|---|---|
| **Intergenic Distal** | - | | | | |
| **Intergenic Proximal** | p = 5.6e-12 | - | | | |
| **Promoter** | p = 2e-16 | p = 4.4e-11 | - | | |
| **Proximal Gene Body** | p = 2e-16 | p = 2e-16 | p = 7.8e-14 | - | |
| **Distal Gene Body** | p = 2e-16 | p = 0.77 | p = 2e-16 | p = 2e-16 | - |

**Supp Fig 15. Pairwise comparisons of changes in H3K27ac at genomic regions using Wilcoxon rank sum test, two sided.**

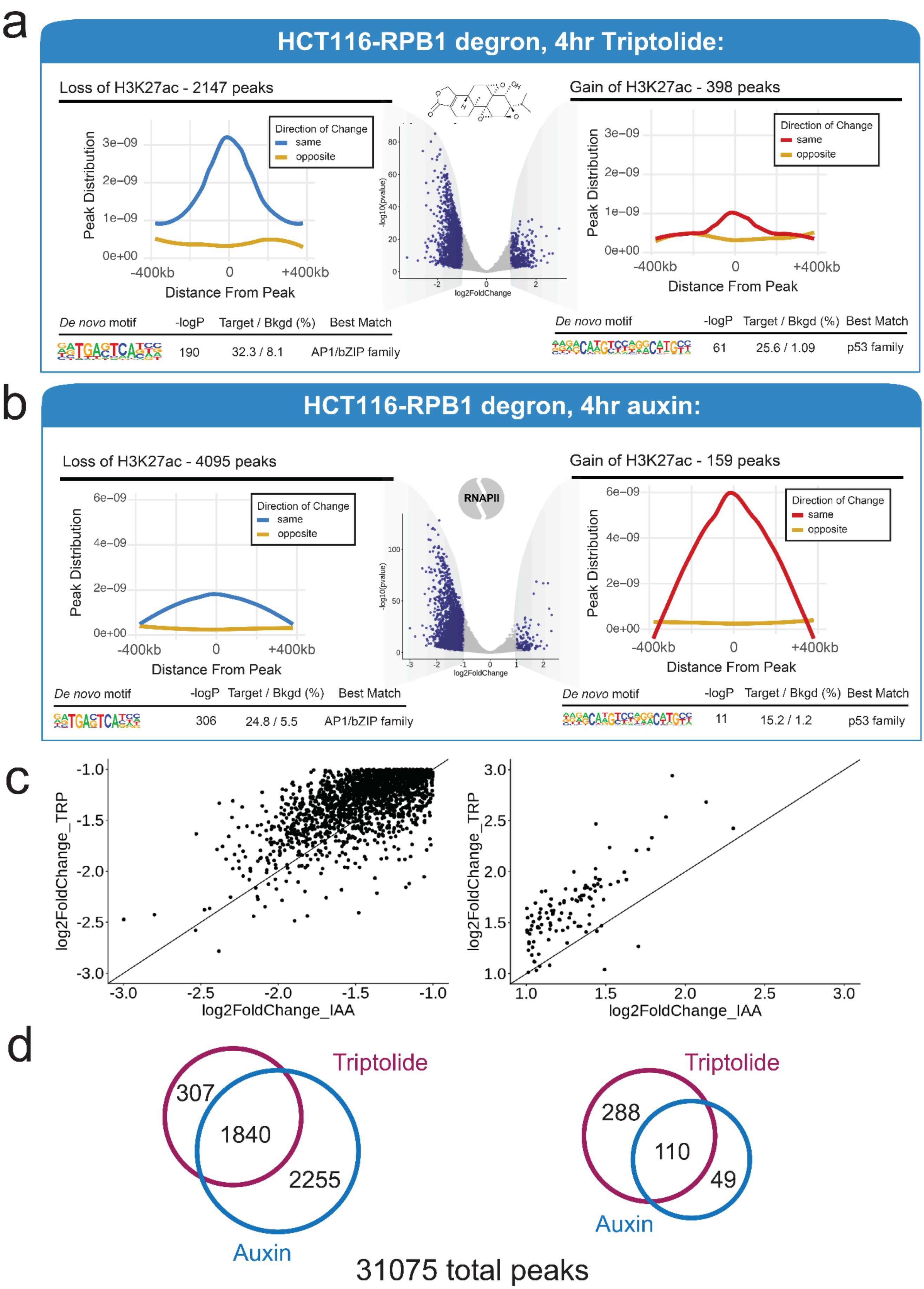
a
HCT116-RPB1 degron, 4hr Triptolide:
Loss of H3K27ac - 2147 peaks
Gain of H3K27ac - 398 peaks
Direction of Change
same
opposite
Peak Distribution
Distance From Peak
-400kb
0
+400kb
log2FoldChange
-log10(pvalue)
De novo motif
-logP
Target / Bkgd (%)
Best Match
190
32.3 / 8.1
AP1/bZIP family
61
25.6 / 1.09
p53 family
b
HCT116-RPB1 degron, 4hr auxin:
Loss of H3K27ac - 4095 peaks
Gain of H3K27ac - 159 peaks
RNAPII
306
24.8 / 5.5
AP1/bZIP family
11
15.2 / 1.2
p53 family
c
log2FoldChange_TRP
log2FoldChange_IAA
d
Triptolide
Auxin
307
1840
2255
288
110
49
31075 total peaks

**Supp. Fig. 16. Differential H3K27ac Peak Finding after Triptolide or auxin-mediated degradation of Rpb1.** Identification of differentially regulated H3K27ac peaks by DESeq2 with custom size factors from ChIP-wrangler.

**a.** Differentially regulated H3K27ac peaks after 4hr Triptolide in HCT116-RPB1 degron cells.

**b.** Differentially regulated H3K27ac peaks after 4hr auxin treatment in HCT116-RPB1 degron cells.

**c.** Correlation of DESeq2, apeglm-shrunken Log2FC in H3K27ac signal between 4 hours of Triptolide and auxin treatments.

**d.** Overlap of differential H3K27ac peaks after 4 hours of Triptolide and auxin treatments, total peaks below.

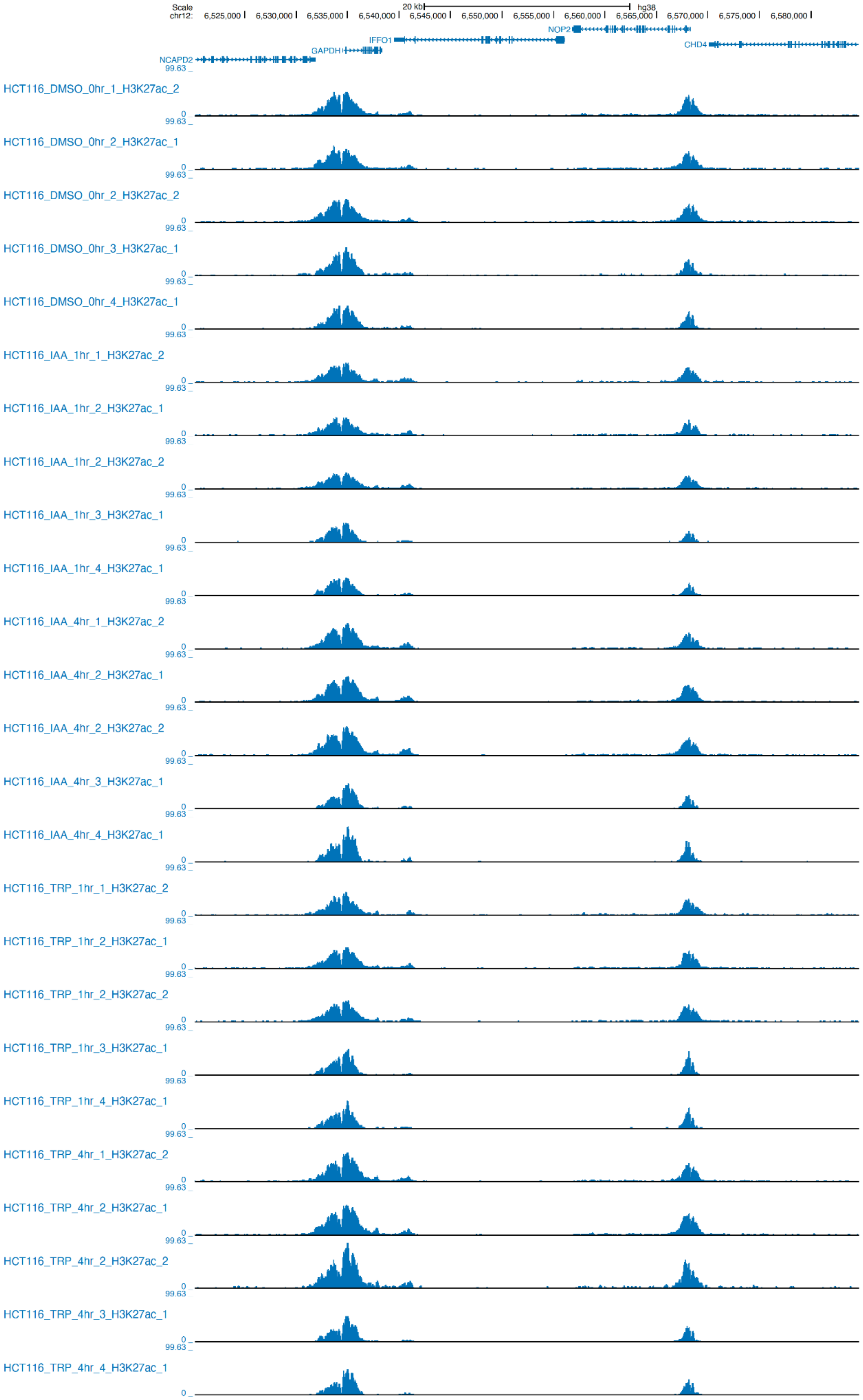
Scale
chr12:
20 kb
hg38
6,525,000
6,530,000
6,535,000
6,540,000
6,545,000
6,550,000
6,555,000
6,560,000
6,565,000
6,570,000
6,575,000
6,580,000
NOP2
IFFO1
CHD4
GAPDH
NCAPD2
99.63
0
HCT116_DMSO_0hr_1_H3K27ac_2
HCT116_DMSO_0hr_2_H3K27ac_1
HCT116_DMSO_0hr_2_H3K27ac_2
HCT116_DMSO_0hr_3_H3K27ac_1
HCT116_DMSO_0hr_4_H3K27ac_1
HCT116_IAA_1hr_1_H3K27ac_2
HCT116_IAA_1hr_2_H3K27ac_1
HCT116_IAA_1hr_2_H3K27ac_2
HCT116_IAA_1hr_3_H3K27ac_1
HCT116_IAA_1hr_4_H3K27ac_1
HCT116_IAA_4hr_1_H3K27ac_2
HCT116_IAA_4hr_2_H3K27ac_1
HCT116_IAA_4hr_2_H3K27ac_2
HCT116_IAA_4hr_3_H3K27ac_1
HCT116_IAA_4hr_4_H3K27ac_1
HCT116_TRP_1hr_1_H3K27ac_2
HCT116_TRP_1hr_2_H3K27ac_1
HCT116_TRP_1hr_2_H3K27ac_2
HCT116_TRP_1hr_3_H3K27ac_1
HCT116_TRP_1hr_4_H3K27ac_1
HCT116_TRP_4hr_1_H3K27ac_2
HCT116_TRP_4hr_2_H3K27ac_1
HCT116_TRP_4hr_2_H3K27ac_2
HCT116_TRP_4hr_3_H3K27ac_1
HCT116_TRP_4hr_4_H3K27ac_1

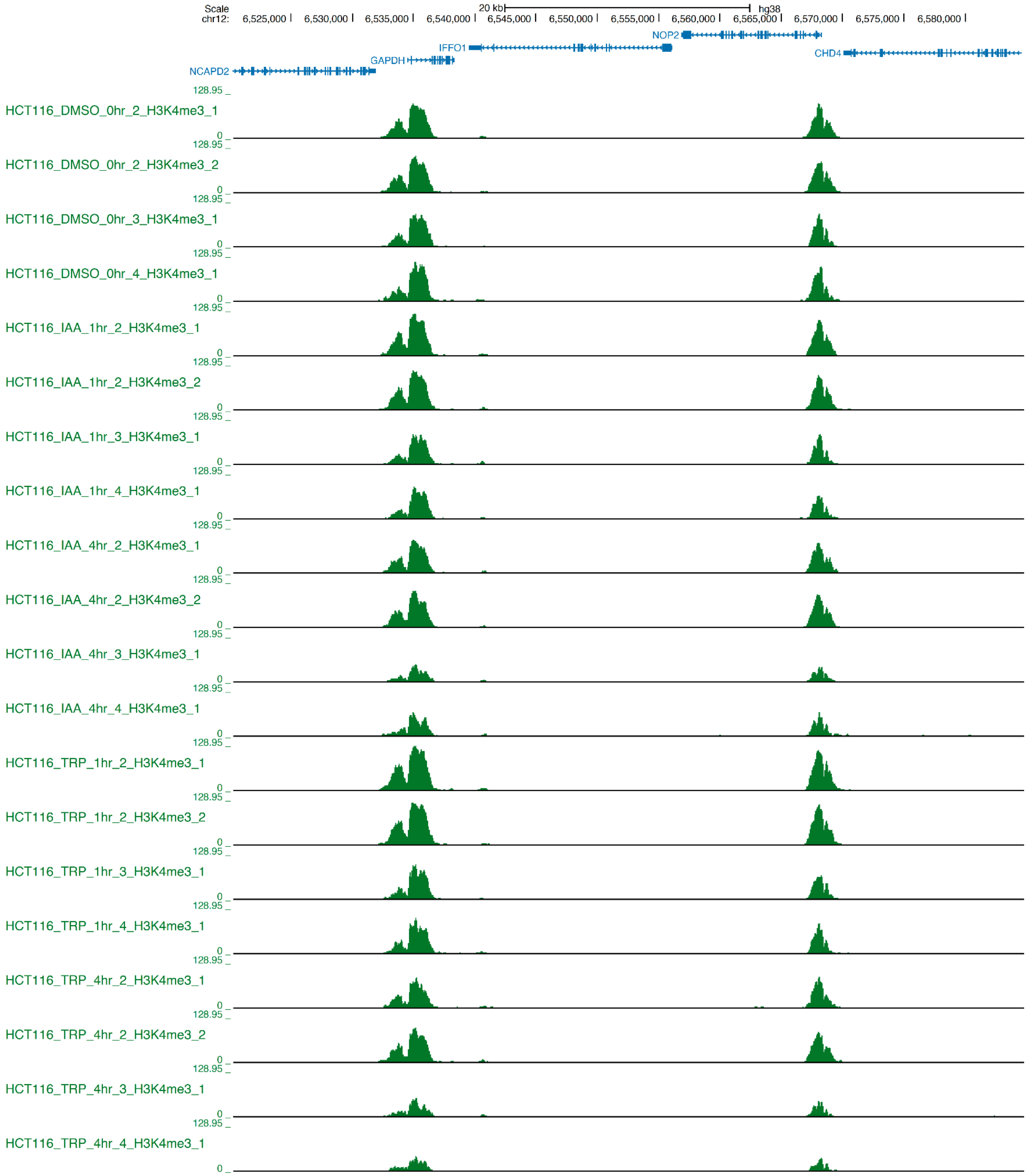

Scale
chr12:
20 kb
hg38
6,525,000
6,530,000
6,535,000
6,540,000
6,545,000
6,550,000
6,555,000
6,560,000
6,565,000
6,570,000
6,575,000
6,580,000
NOP2
IFFO1
CHD4
GAPDH
NCAPD2
128.95
0
HCT116_DMSO_0hr_2_H3K4me3_1
HCT116_DMSO_0hr_2_H3K4me3_2
HCT116_DMSO_0hr_3_H3K4me3_1
HCT116_DMSO_0hr_4_H3K4me3_1
HCT116_IAA_1hr_2_H3K4me3_1
HCT116_IAA_1hr_2_H3K4me3_2
HCT116_IAA_1hr_3_H3K4me3_1
HCT116_IAA_1hr_4_H3K4me3_1
HCT116_IAA_4hr_2_H3K4me3_1
HCT116_IAA_4hr_2_H3K4me3_2
HCT116_IAA_4hr_3_H3K4me3_1
HCT116_IAA_4hr_4_H3K4me3_1
HCT116_TRP_1hr_2_H3K4me3_1
HCT116_TRP_1hr_2_H3K4me3_2
HCT116_TRP_1hr_3_H3K4me3_1
HCT116_TRP_1hr_4_H3K4me3_1
HCT116_TRP_4hr_2_H3K4me3_1
HCT116_TRP_4hr_2_H3K4me3_2
HCT116_TRP_4hr_3_H3K4me3_1
HCT116_TRP_4hr_4_H3K4me3_1

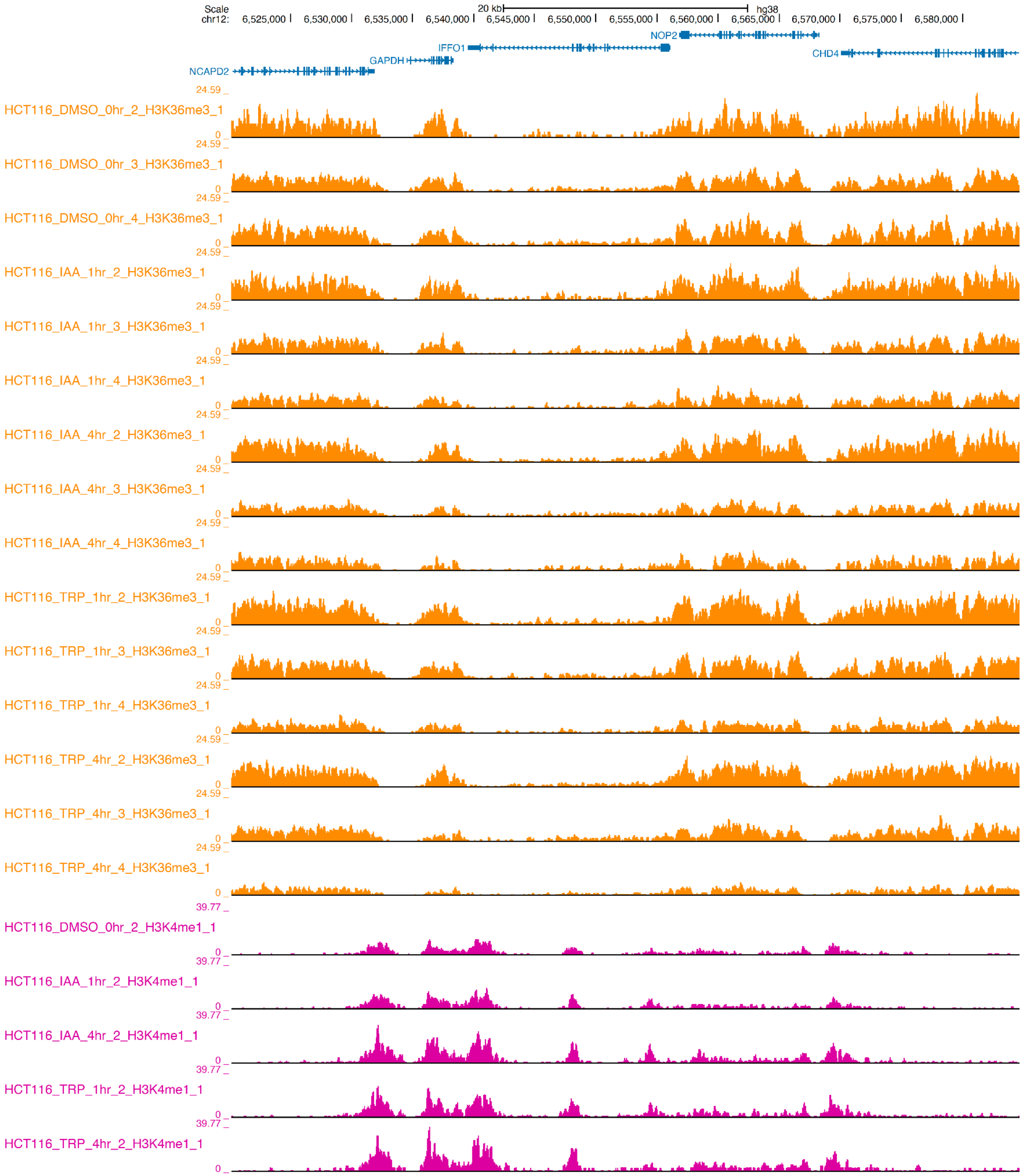
Scale
chr12:
20 kb
hg38
6,525,000
6,530,000
6,535,000
6,540,000
6,545,000
6,550,000
6,555,000
6,560,000
6,565,000
6,570,000
6,575,000
6,580,000
NOP2
IFFO1
CHD4
GAPDH
NCAPD2
24.59
0
HCT116_DMSO_0hr_2_H3K36me3_1
HCT116_DMSO_0hr_3_H3K36me3_1
HCT116_DMSO_0hr_4_H3K36me3_1
HCT116_IAA_1hr_2_H3K36me3_1
HCT116_IAA_1hr_3_H3K36me3_1
HCT116_IAA_1hr_4_H3K36me3_1
HCT116_IAA_4hr_2_H3K36me3_1
HCT116_IAA_4hr_3_H3K36me3_1
HCT116_IAA_4hr_4_H3K36me3_1
HCT116_TRP_1hr_2_H3K36me3_1
HCT116_TRP_1hr_3_H3K36me3_1
HCT116_TRP_1hr_4_H3K36me3_1
HCT116_TRP_4hr_2_H3K36me3_1
HCT116_TRP_4hr_3_H3K36me3_1
HCT116_TRP_4hr_4_H3K36me3_1
39.77
HCT116_DMSO_0hr_2_H3K4me1_1
HCT116_IAA_1hr_2_H3K4me1_1
HCT116_IAA_4hr_2_H3K4me1_1
HCT116_TRP_1hr_2_H3K4me1_1
HCT116_TRP_4hr_2_H3K4me1_1

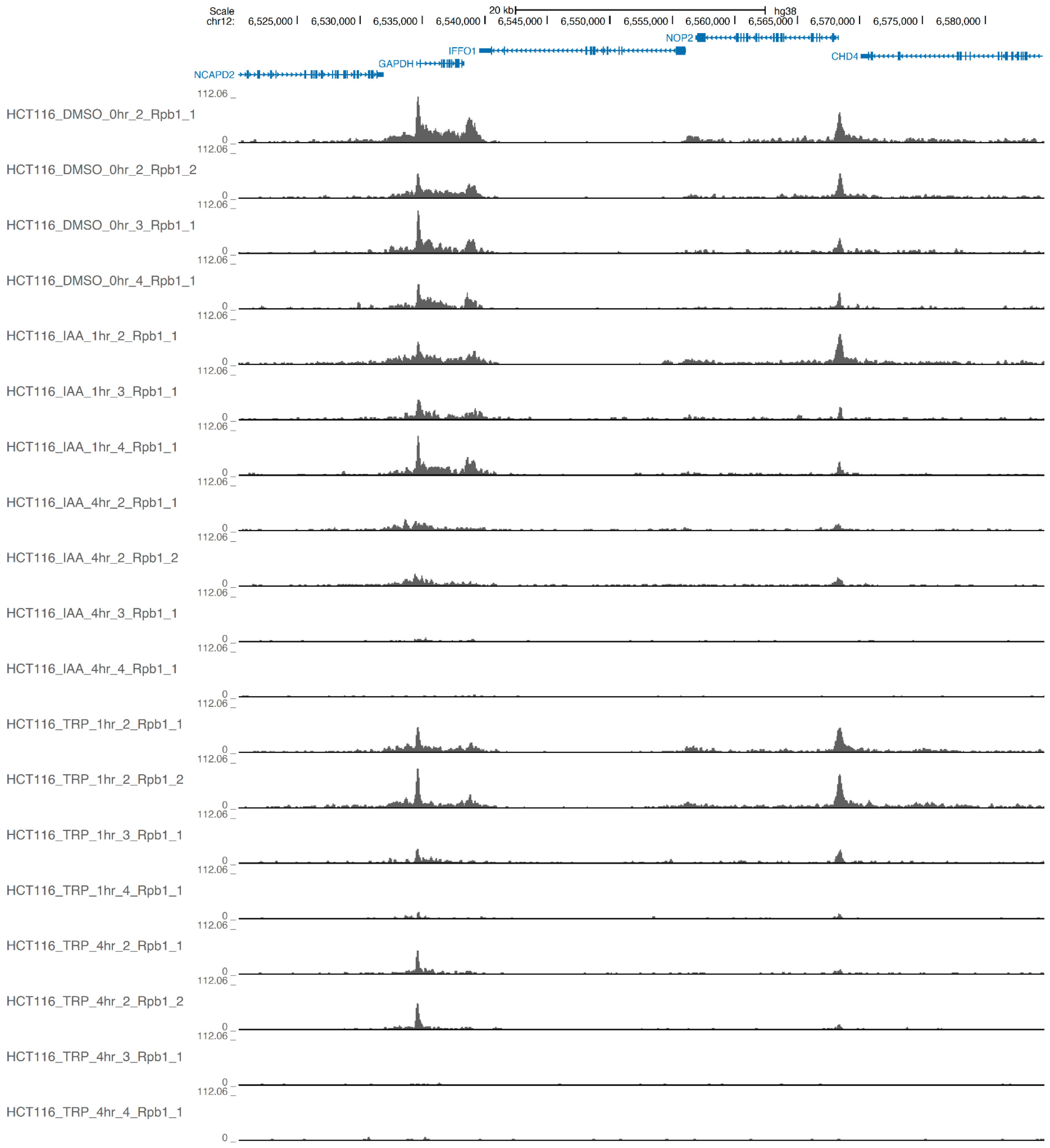


**Supp Fig. 17. UCSC genome browser tracks for additional biological replicates of treatment of HCT116-RPB1 degron cells with Triptolide an auxin.**

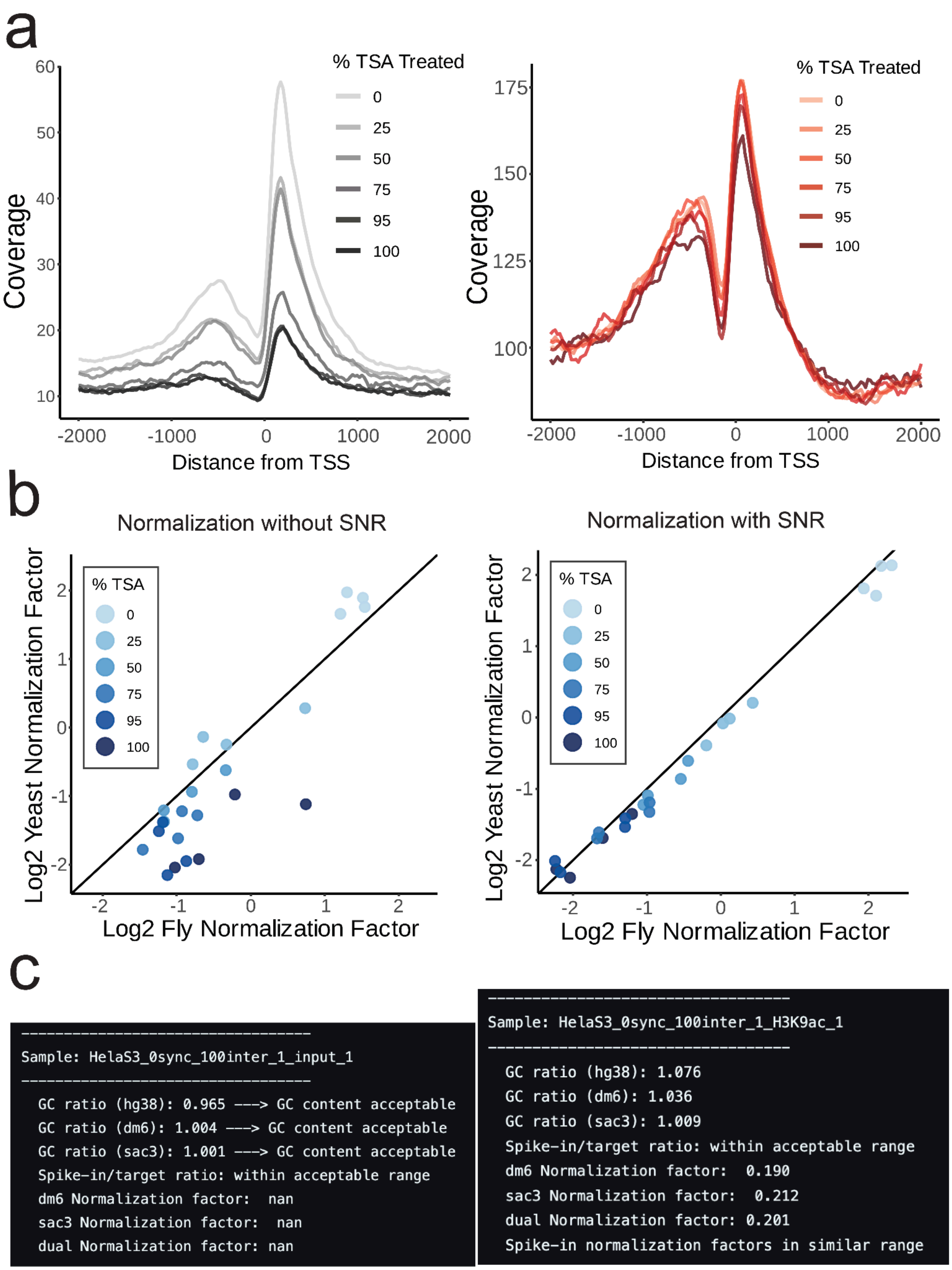


**Supp. Fig. 18. Examples of expected quality metric results provided by ChIP-wrangler.**
**a.** Histograms of H3K27ac ChIP-seq signal at RefSeq TSSs in *D. melanogaster* and *S. cerevisiae*.
**b.** Correlation of normalization factors derived from *D. melanogaster* and *S. cerevisiae*, before and after accounting for variations in IP efficiency with spike-ins.
**c.** Examples of ChIP-wrangler output log file reports for input (left) and IP samples (right); quality metrics such as spike-in/target ratio, GC content relative to expected, and correlation of normalization factors are shown.
