## Supplementary material for "Improved spike-in normalization clarifies the relationship between active histone modifications and transcription": patel-etal-biorxiv-sup-material-revision2-submit

### IAA treatment

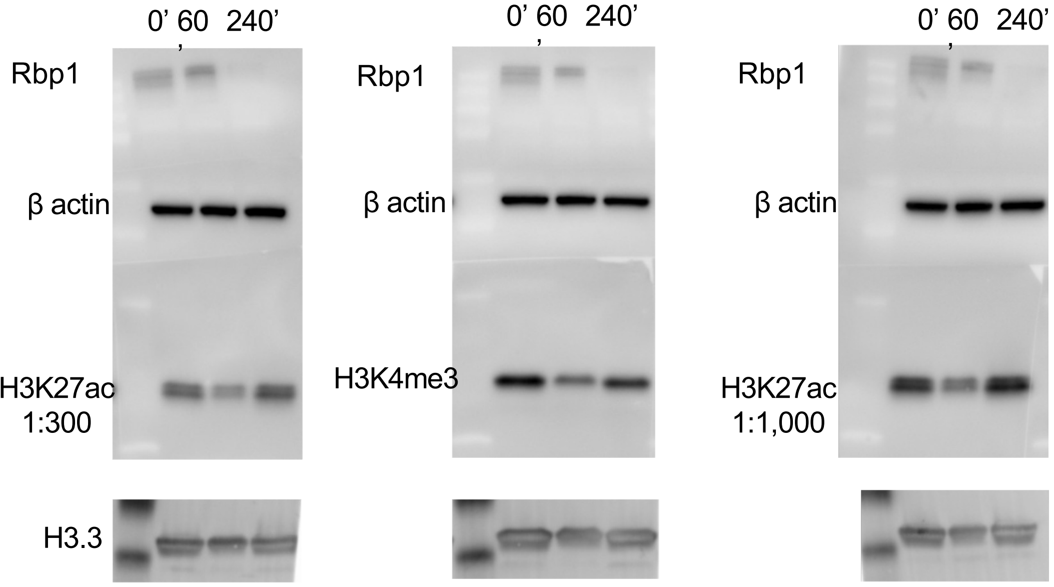


10.25.2024

### IAA treatment

12.6 μg protein

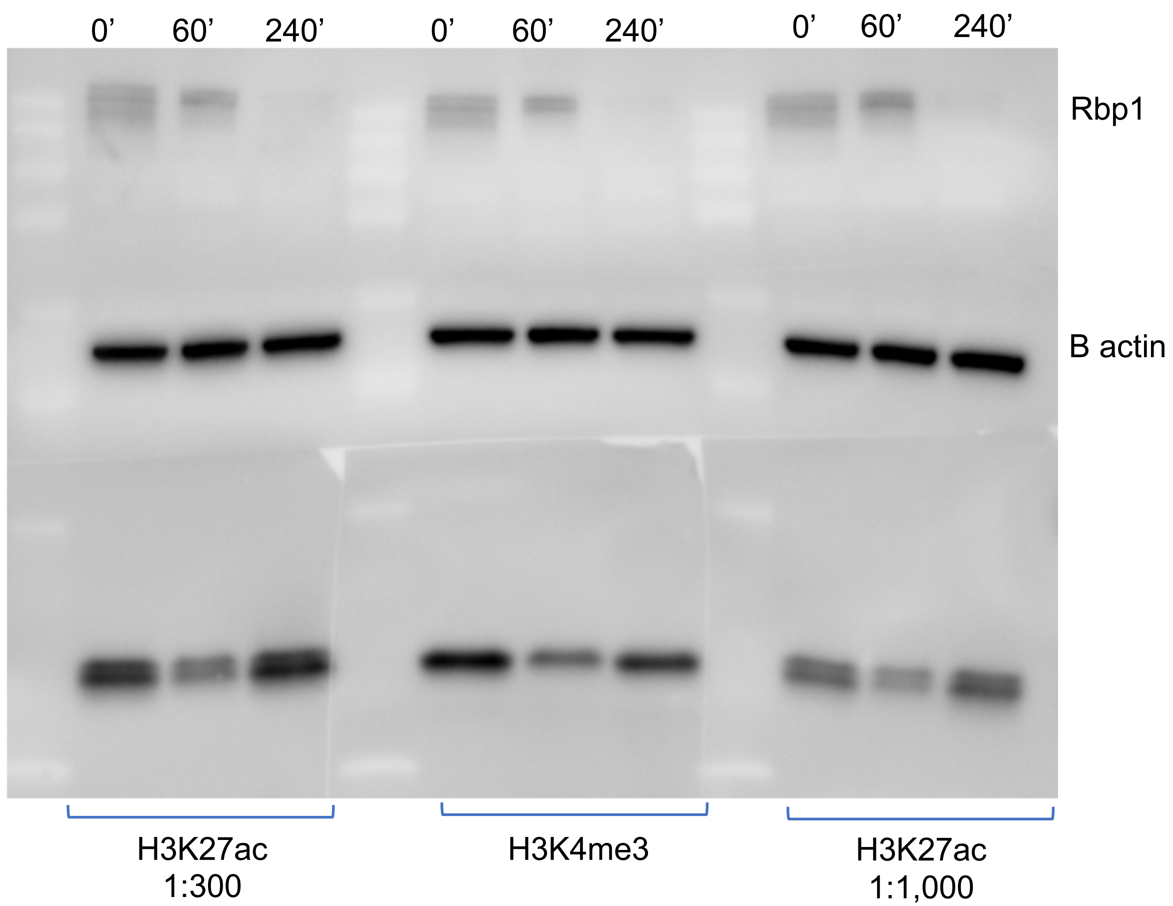

Trp treatment

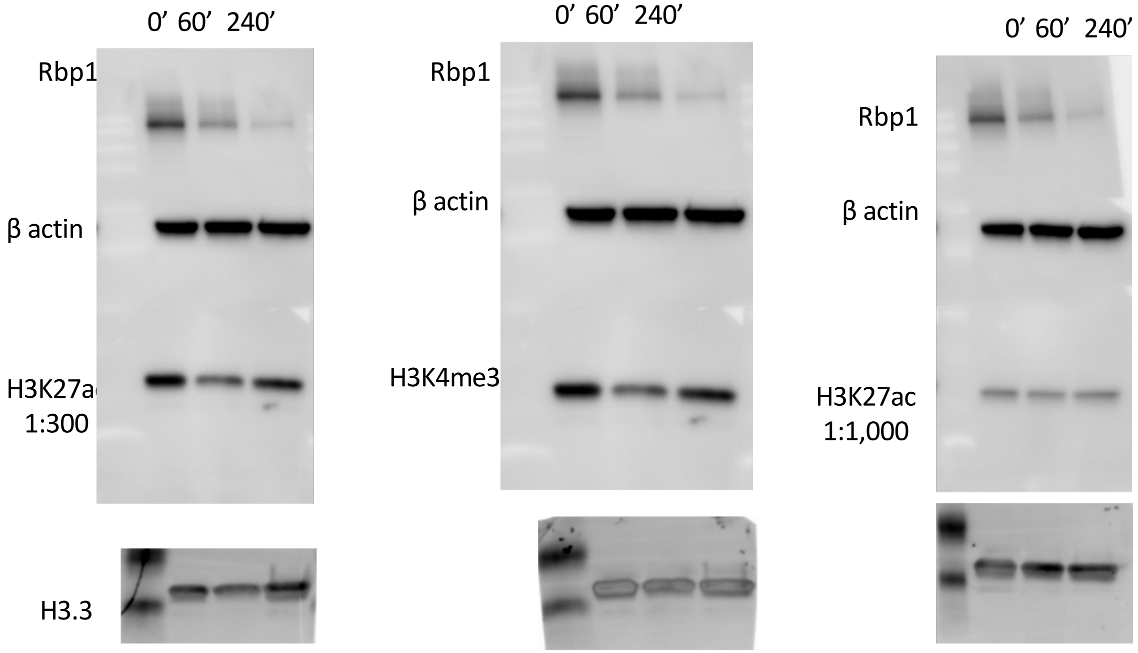


Trp treatment

12.6 µg protein

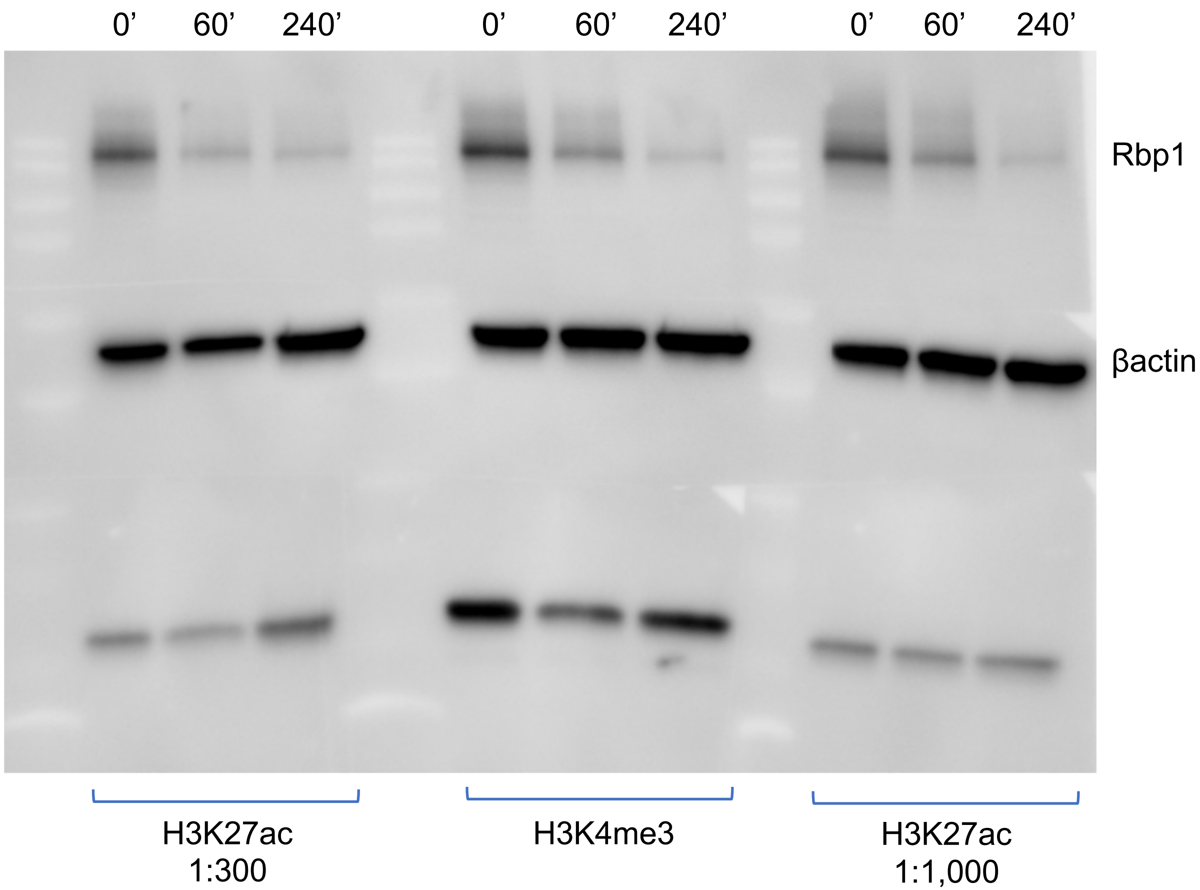

11.1.2024

IAA treatment

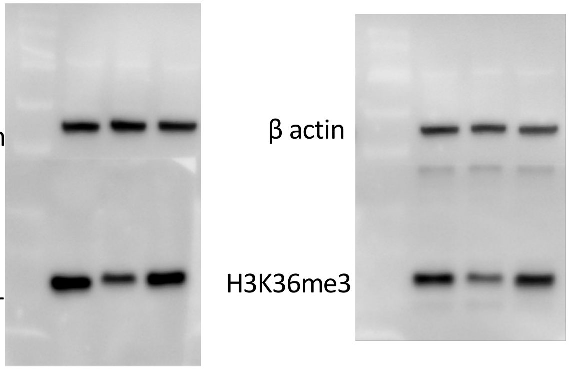


Trp treatment

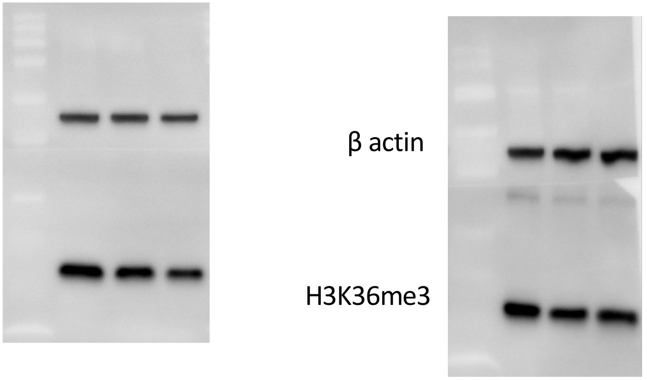


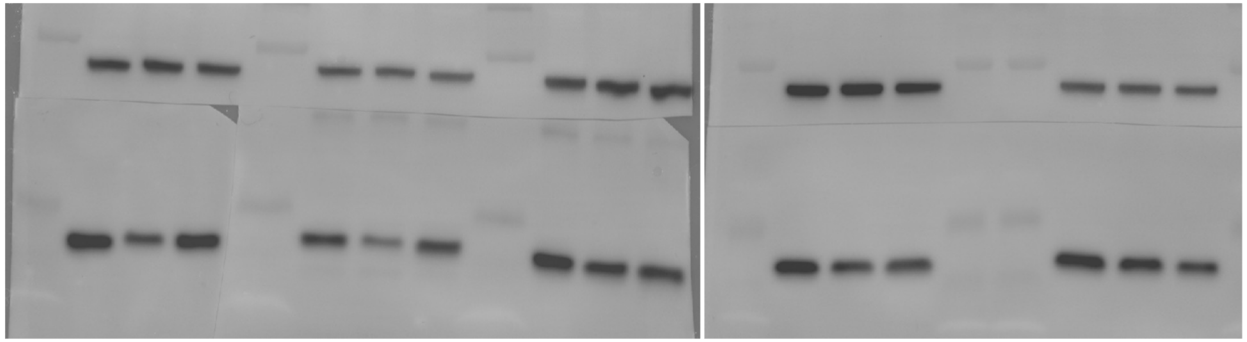


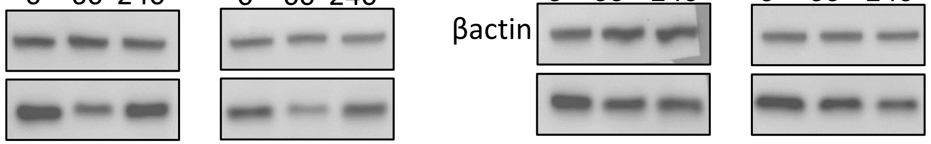

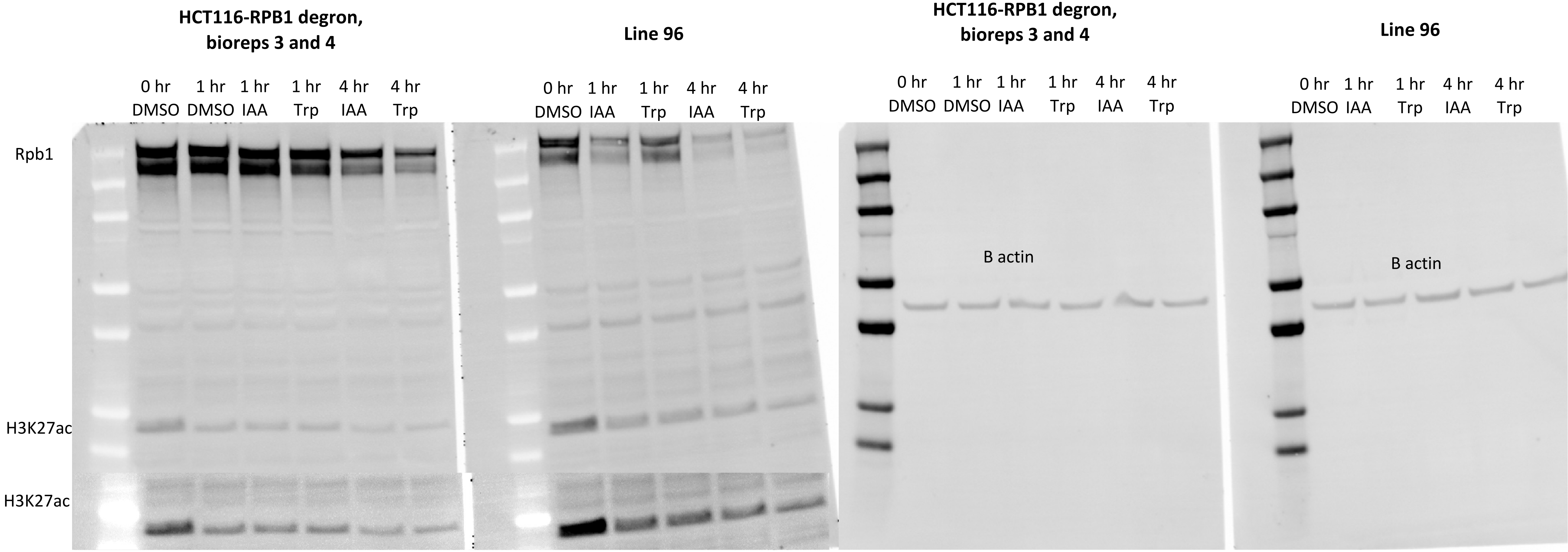
HCT116-RPB1 degron, bioreps 3 and 4
Line 96
HCT116-RPB1 degron, bioreps 3 and 4
Line 96
0 hr DMSO
1 hr DMSO
1 hr IAA
1 hr Trp
4 hr IAA
4 hr Trp
0 hr DMSO
1 hr IAA
1 hr Trp
4 hr IAA
4 hr Trp
0 hr DMSO
1 hr DMSO
1 hr IAA
1 hr Trp
4 hr IAA
4 hr Trp
0 hr DMSO
1 hr IAA
1 hr Trp
4 hr IAA
4 hr Trp
Rpb1
H3K27ac
H3K27ac
B actin
B actin

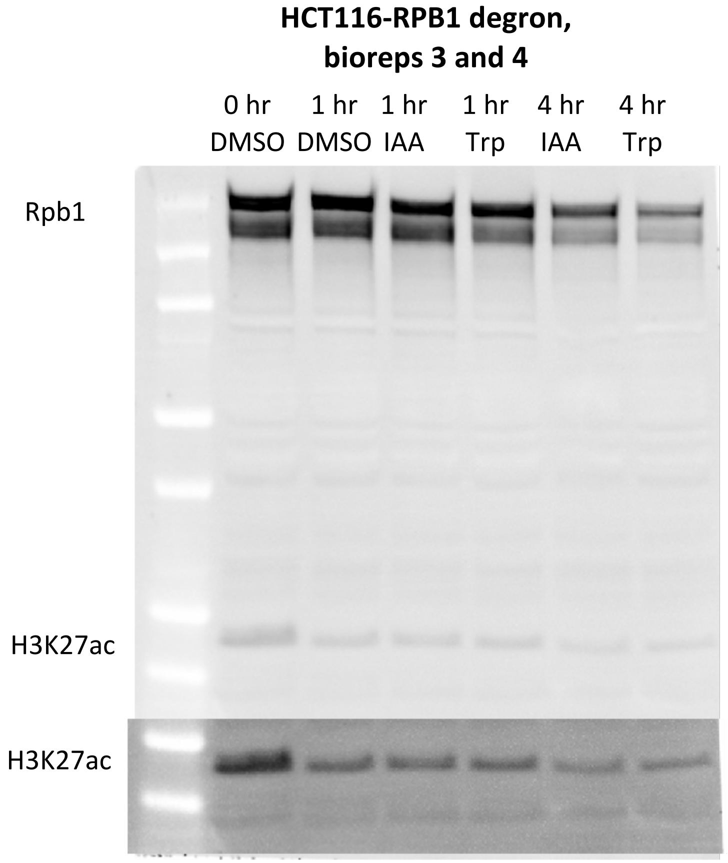
HCT116-RPB1 degron,
bioreps 3 and 4
0 hr DMSO
1 hr DMSO
1 hr IAA
1 hr Trp
4 hr IAA
4 hr Trp
Rpb1
H3K27ac
H3K27ac

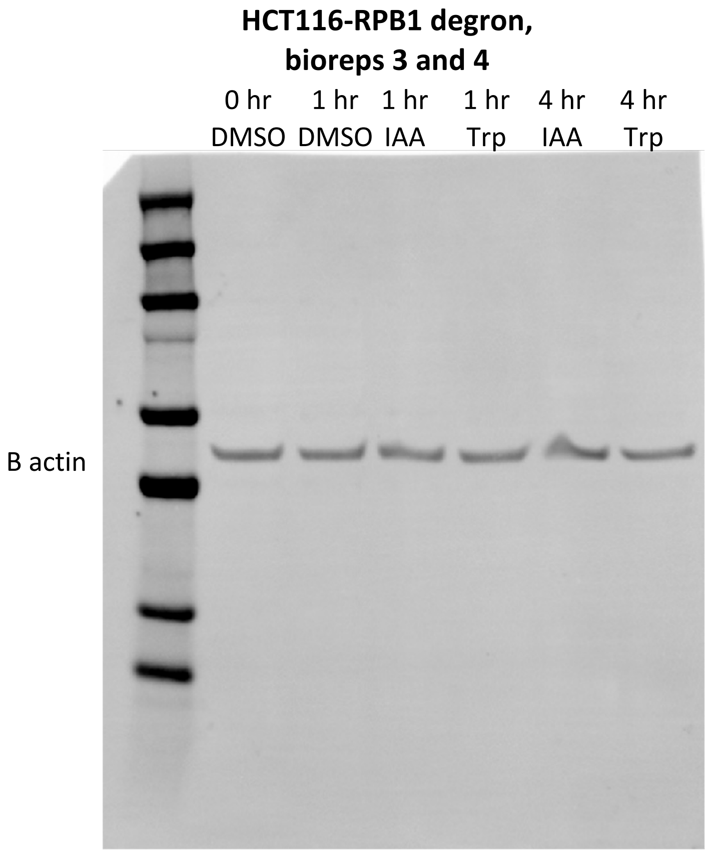
HCT116-RPB1 degron,
bioreps 3 and 4
0 hr DMSO
1 hr DMSO
1 hr IAA
1 hr Trp
4 hr IAA
4 hr Trp
B actin

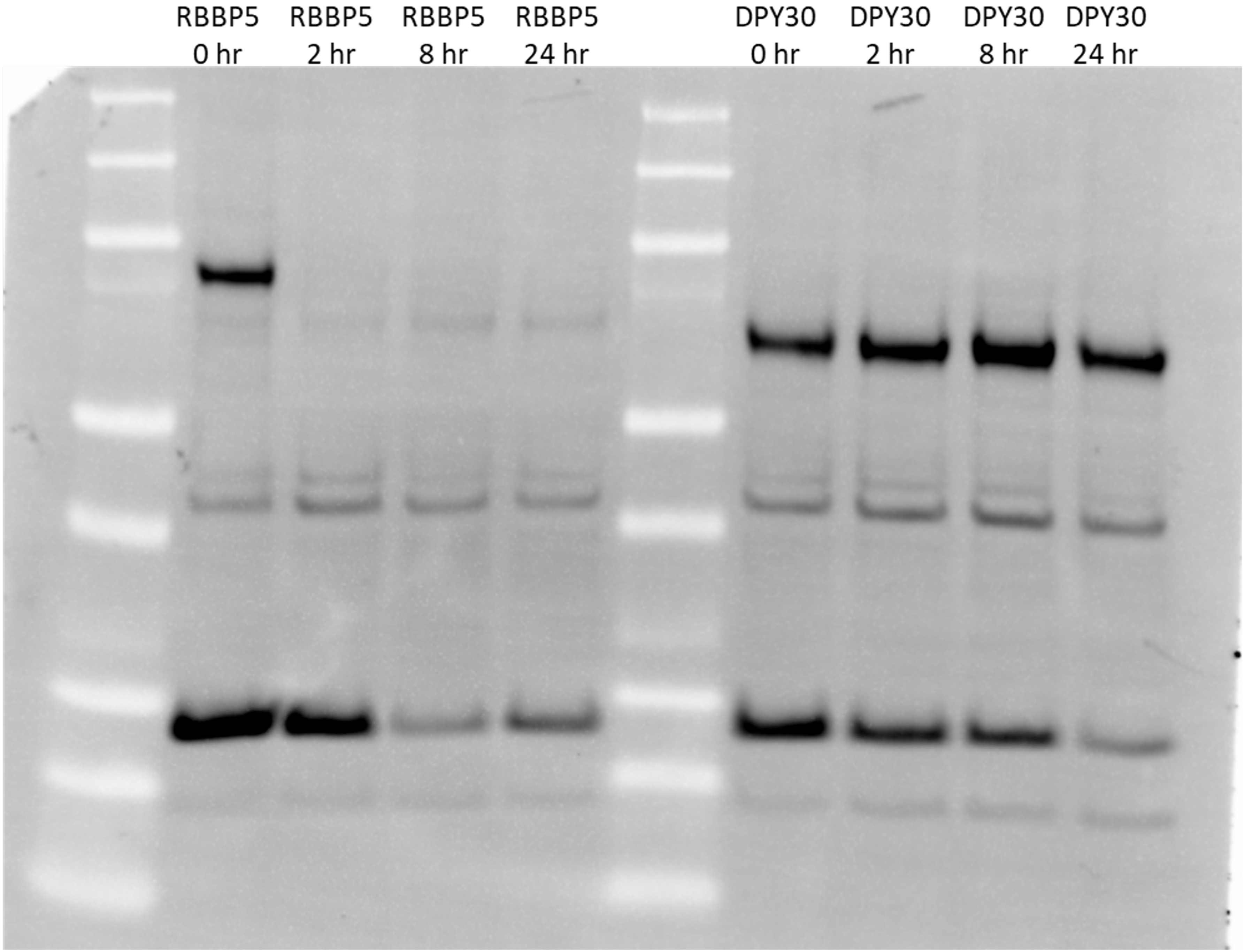
RBBP5 0 hr
RBBP5 2 hr
RBBP5 8 hr
RBBP5 24 hr
DPY30 0 hr
DPY30 2 hr
DPY30 8 hr
DPY30 24 hr
